# Biological Sex Differences after High and Low Doses of Influenza A Virus Infection in a Diet-induced Obesity Mouse Model

**DOI:** 10.1101/2025.11.18.689021

**Authors:** Saurav Pantha, Brian Wolfe, Saranya Vijayakumar, Shristy Budha Magar, Tawfik Aboellail, Santosh Dhakal

## Abstract

Obesity is an independent risk factor for influenza A virus (IAV) pathogenesis. In non-obese hosts, following IAV infection, adult females develop more severe disease than males. Limited research on biological sex differences following IAV infection in individuals with obesity forms the basis for this study. Male and female C57BL/6J mice were treated with high-fat or low-fat diets to obtain mice with or without obesity. They were inoculated intranasally with a high (10^3^) or low dose (10^1.5^ TCID_50_) of mouse-adapted A/California/04/2009 H1N1 IAV and followed for 21 days post-infection (dpi) for morbidity. Subsets of mice were euthanized at different dpi to measure lung virus titers, histopathological changes, cellular infiltration, and cytokine induction. After a high-dose infection, females with obesity demonstrated shorter median survival time, greater parenchymal inflammation, and heightened induction of inflammatory markers than males with obesity. After a low-dose infection, females than males, and mice with obesity than the non-obese mice, exhibited severe morbidity. Female mice with obesity exhibited the greatest disease severity, where 25% (2/8) reached humane endpoint. Delayed but persistent inflammatory changes were observed in females with obesity, characterized by relatively lower pathological changes, lesser induction of cytokines and chemokines, and reduced myeloid and lymphoid infiltration in the lungs at 3 dpi followed by sustained pathological changes and cytokine and chemokine induction at 21 dpi. Our study suggests that the biological sex difference following IAV infection persists during obesity. Females with obesity experience greater influenza disease severity than males, possibly driven by more severe dysregulation of inflammatory responses.

**Importance:** The prevalence of obesity is increasing worldwide. The risk of developing severe disease following influenza virus infection is higher in individuals with obesity. Males and females with obesity differ in terms of fat deposition and inflammatory and metabolic biomarkers. These differences can alter the outcomes of influenza-induced disease pathogenesis. The observation of sex differences in IAV pathogenesis among non-obese hosts underscores the importance of determining whether similar differences occur in the context of obesity. Indeed, our study suggested that females with obesity experience greater disease severity than males with obesity after IAV infection. Understanding the immunological mechanisms of such differences will advance precision medicine approaches to develop safer and more effective preventive and therapeutic strategies against influenza for the high-risk populations with obesity.

## Introduction

Obesity (i.e., having a body mass index, BMI ≥30) is characterized by excessive fat deposition and a state of chronic low-grade inflammation in the body. Since the 1990s, obesity has doubled in adults and quadrupled among children and adolescents, and more than 1 billion people in the world are living with obesity (1). By 2035, 51% of the world’s population is estimated to be either overweight or obese (2). In the US, 2 in 5 adults were living with obesity in 2023, and it is estimated that by 2050, at least 1 in 5 children (5-14 years), 1 in 3 adolescents (15-24 years), and 2 in 3 adults (≥25 years) will have obesity (3, 4). Dietary intake, such as increased consumption of ultra-processed foods, is a major factor driving the global rise in obesity (5).

Influenza viruses cause over a billion cases of respiratory illnesses annually, with 3-5 million severe cases and up to 650,000 deaths (6). Obesity was recognized as a risk factor for severe outcomes of influenza, including increased hospitalizations, the need for intensive care, and deaths during the 2009 H1N1 influenza A virus (IAV) pandemic (7). This has been observed subsequently, even during seasonal influenza outbreaks, further confirming obesity as an independent risk factor for influenza-associated disease severity (8, 9). A systematic review and meta-analysis showed that people with obesity have a 56% and 100% higher risk of severe disease and deaths associated with influenza, respectively, compared to people without obesity (8). Symptomatic adults with obesity are likely to shed IAV 42% longer than non-obese adults, while asymptomatic adults with obesity shed IAV 104% longer (10). Human and animal model studies have shown that obesity is linked to delayed virus clearance, progression to viral pneumonia, emergence of IAV strains, and a higher risk of secondary bacterial infection (11–15).

Biological sex (i.e., being male or female based on chromosomal and hormonal differences) has an important role in IAV pathogenesis and immunity (16, 17). Epidemiological data in humans show that adult females of reproductive age are more likely to experience higher influenza-related hospitalization rates than their age-matched males (18, 19). Analysis of age-standardized notification rates of laboratory-confirmed cases of influenza between 2009 and 2015 in Australia showed significantly higher cases in adult females for both the H1N1 and H3N2 IAVs compared to adult males, suggesting that females are more susceptible to IAVs (20). The IAV challenge study in humans also showed that adult females a had more symptoms and a longer duration of symptoms than males (21). The non-obese mouse model of IAV infection shows that females suffer a worse outcome from IAV infection than males, associated with elevated inflammatory immune responses and pulmonary tissue damage, but comparable pulmonary virus replication (22, 23). In mouse models, elevated levels of testosterone and amphiregulin (i.e., a protein that drives lung tissue repair) in adult males were associated with better influenza outcomes in males, while administration of high doses of estradiol reduced the severe outcomes in females (22, 23), indicating the roles of sex steroids in mediating sex differences in non-obese hosts.

Sex difference exists in adiposity in humans, where females tend to have more subcutaneous adipose tissue and males tend to have more visceral adipose tissue accumulation (24). Both subcutaneous and visceral adipose tissues have receptors for sex steroids and are involved in mediating inflammatory responses (24, 25). Males and females with obesity also differ in terms of adipokine and inflammatory cytokine concentrations (26). Though sex differences during IAV infection are studied in non-obese hosts, such studies in hosts with diet-induced obesity (DIO) are limited. In a prior study, Alarcon et al. used a mouse model of DIO and IAV infection. They observed that obesity-associated effects on lung inflammation, cytokine and chemokine production, and immune cell infiltration were comparable between males and females (27). However, that study only used a lethal dose of IAV; analysis was performed only at 5 days post-infection (dpi) and lacked temporal analysis (27). In this study, we used a DIO mouse model to investigate biological sex differences in virus replication, pulmonary inflammation, and disease severity following infection with high- and low-dose 2009 pandemic H1N1 IAV infection. IAV-infected mice were followed for morbidity measurements up to 21 dpi, with subsets of mice euthanized at 3, 10, and 21 dpi to compare temporal changes in influenza pathogenesis. Our data suggests that biological sex impacts IAV pathogenesis, where females with obesity suffer from more severe disease compared to males.

## Results

### Males and females differed in metabolic biomarkers and in the progression of obesity

Before the diet treatment (i.e., week 0), there was no difference in body mass between males on LFD and HFD, or between females on LFD and HFD (**Supplementary Figure 3A**). Males, however, had significantly greater body mass than the females. After the diet treatment, the percentage body mass change in male mice on HFD was significantly greater than their LFD controls within 1^st^ week, while it was substantially greater in females only from the 6^th^ week (**Supplementary Figure 3B**). While 100% of males on HFD became obese within the 9^th^ week, only 70% of females on HFD became obese by the 14^th^ week, and 30% did not reach the obesity definition and were defined as non-responders (**Supplementary Figure 3C-D**). After 14 weeks, both male mice with obesity (O-M) and female mice with obesity (O-F) had significantly greater body mass, BMI, and deposition of gonadal, inguinal, and perirenal adipose tissues compared to non-obese males (NO-M) and non-obese females (NO-F), respectively (**Figure 1A-E**). Even after the 14 weeks of diet treatment, males in each group were bigger in size than the female mice. GTT on the 14^th^ week showed comparable fed glucose levels (**Figure 1F**). Male mice with obesity had significantly higher blood glucose levels at 30, 60, and 120 minutes, while female mice with obesity maintained such a difference at 15, 30, and 60 minutes; and both males and females with obesity had significantly higher GTT AUC values compared to their non-obese controls (**Figure 1G-H**). GTT on the 8^th^ week also showed similar responses in mice that eventually became obese (**Supplementary Figure 3E-G**).

**Figure 1.**
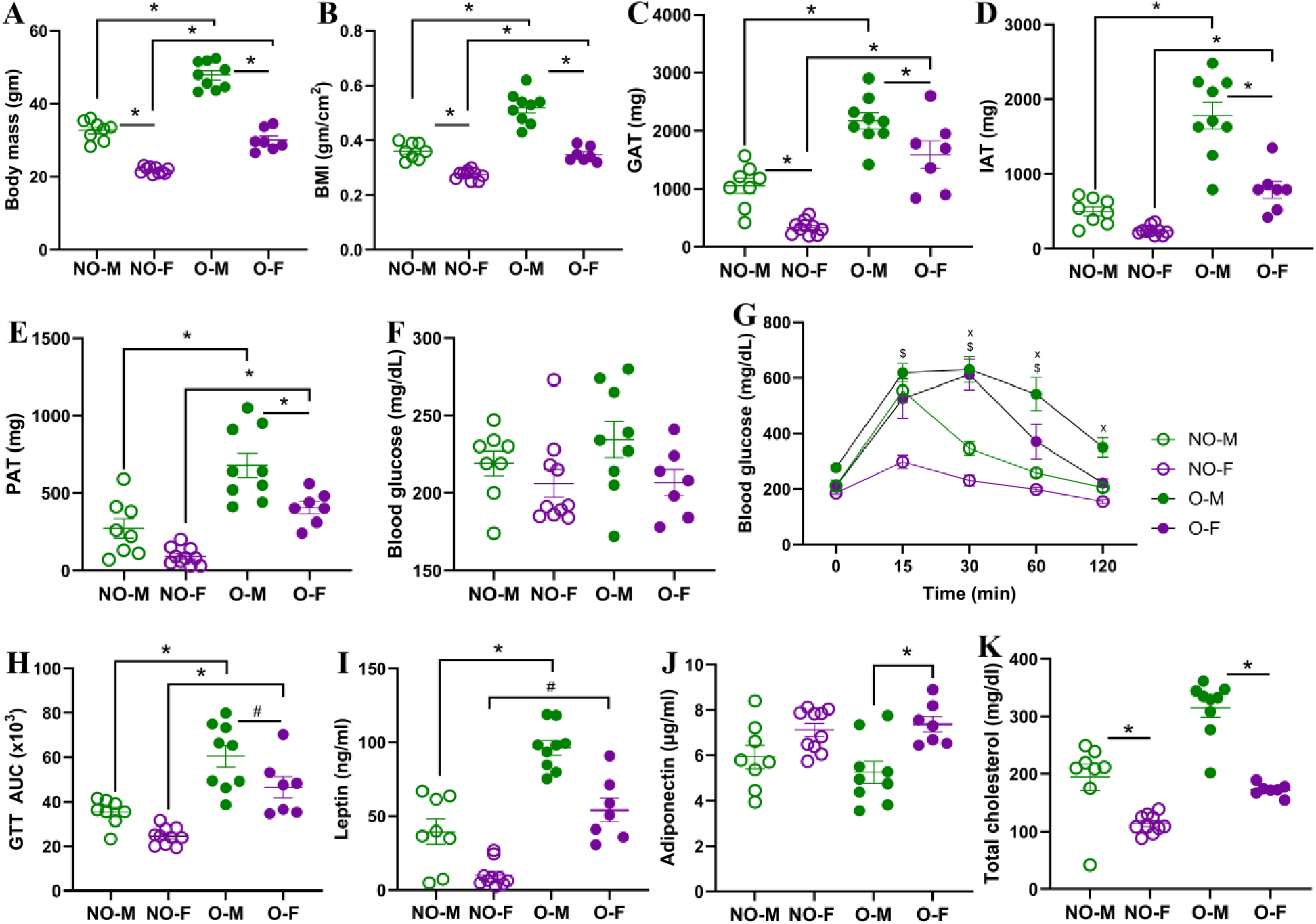
Progression of obesity and metabolic states in male and female mice. Male and female mice were fed with a low-fat diet (LFD) or a high-fat diet (HFD) for 14 weeks. Comparisons of (A) body mass after the 14^th^ week, (B) body mass index (BMI) on the day of infection, and (C) gonadal adipose tissue (GAT), (D) inguinal adipose tissue (IAT), and (E) perirenal adipose tissue (PAT) on day of euthanization (i.e., 3 dpi) are compared between male and female mice, with or without obesity. A glucose tolerance test (GTT) was performed on the 14^th^ week of diet treatment. Comparisons of (F) fed glucose, (G) GTT over time, and (H) GTT area under the curve (AUC) are shown. Comparisons of (I) leptin, (J) adiponectin, and (K) total cholesterol in plasma samples from blood collected at the 14^th^ week are shown. Data is presented as mean ± standard error of the mean (SEM) (n=7-10/group). Statistical comparisons were done using one-way ANOVA followed by Tukey’s post-hoc comparisons or Kruskal-Wallis test followed by Dunn’s post-hoc comparisons. Repeated measurements in 1G were compared by repeated measures ANOVA (mixed model) and Tukey’s multiple comparisons. An asterisk (*) indicates a statistically significant difference (p<0.05) while a hash (^#^) indicates a trend (0.05≤p≤0.1). In 1G, x’ and ‘$’ represent significant differences between NO-M vs O-M, and NO-F vs O-F, respectively. Abbreviations: NO-M, non-obese males; NO-F, non-obese females; O-M, obese males; and O-F, obese females.

At week 0, female mice had significantly higher plasma concentrations of leptin and adiponectin and lower concentrations of total cholesterol than males (**Supplementary Figure 3H-J**). However, after the 14^th^ week of diet treatment, leptin concentration was significantly higher in males with obesity and showed a higher trend in females with obesity compared to their non-obese controls **(Figure 1I)**. At 14 weeks, adiponectin concentration was significantly higher in females with obesity compared to males with obesity **(Figure 1J)**. Total cholesterol at 14 weeks was significantly higher in males than in females, both with and without obesity. Mice with obesity had higher concentrations of total cholesterol in the 14^th^ week (i.e., average concentration in NO-M= 194.5 mg/dL, O-M= 315.2 mg/dL, NO-F= 113.7 mg/dL, and O-F= 172.8 mg/dL), but data were not statistically significant **(Figure 1K)**. When we compared the fold increase in these biomarkers from week 0 to the 14^th^ week of diet treatment, males with obesity had an increased fold change of leptin (p=0.09) and significantly higher total cholesterol (p<0.05), while females with obesity had a higher trend of total cholesterol compared to their respective non-obese controls (**Supplementary Figures 3K-M**). Overall, these data indicate that the progression of obesity is quicker and severe in males than in females on HFD, and males and females with obesity differ in terms of metabolic biomarkers. The responder females (i.e., the obese with ≥20% body mass gain) also showed significantly increased body mass, BMI, adipose tissue deposition, glucose intolerance, and higher leptin and total cholesterol levels than the non-obese females, as observed between males, confirming obesity status.

### After a high-dose influenza virus infection, the median survival time was shorter for females, and lung parenchymal inflammation was greater in female mice with obesity

After infection with the high dose (i.e. 10^3^ TCID_50_) of the 2009 pandemic H1N1 IAV, all virus-infected mice started losing body mass gradually and reached the humane endpoint (i.e., ≥25%) except for one male with obesity which lost up to 21% body mass and even at 21 dpi it lost 17% (**Figure 2A**). Likewise, body temperature decreased in infected mice up to 10 dpi or their humane endpoint (**Figure 2B**). Sex difference was observed in survival as the median survival time was 8 days for non-obese females, 12 days for non-obese males, 9 days for females with obesity, and 13 days for males with obesity (**Figure 2C**). At 3 dpi, there was no difference in infectious virus titers in the lungs by biological sex or obesity status of the mice, indicating that sex difference in disease severity was not attributed to differential pulmonary virus replication (**Figure 2D**).

**Figure 2.**
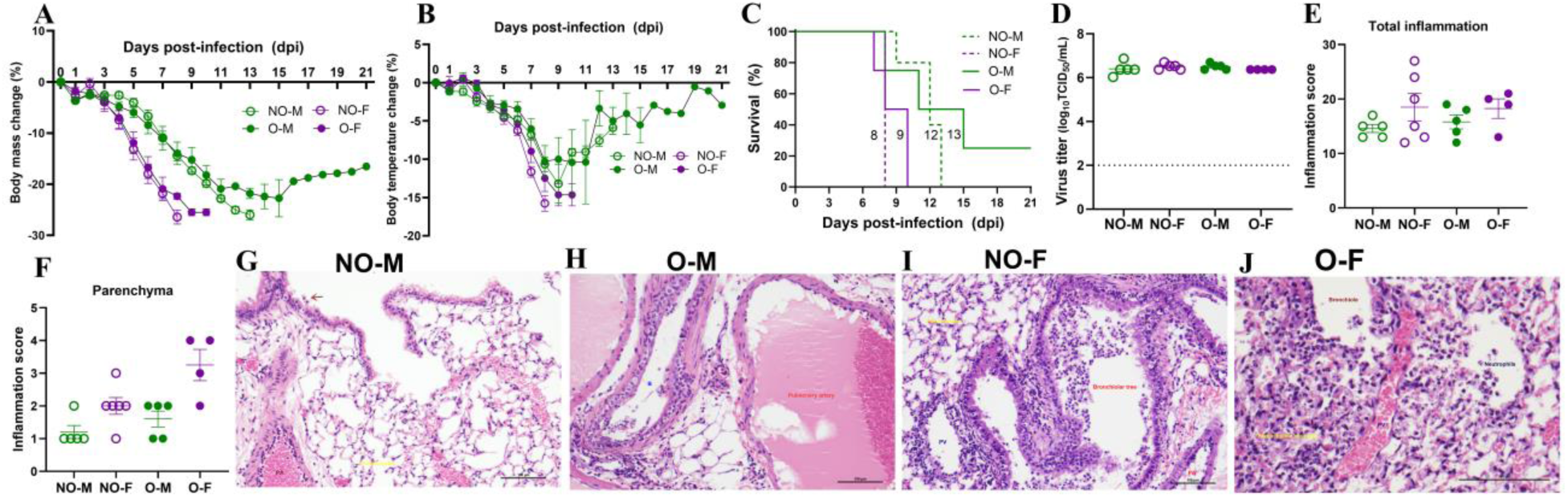
Morbidity, virus replication, and lung histopathology following a high-dose IAV infection. Male and female mice, with or without obesity, were intranasally inoculated with a high dose (10^3^ TCID_50_) of the 2009 pandemic H1N1 IAV or vehicle (i.e., medium) only and monitored for body mass and body temperature changes. (A) Percentage change in body mass and (B) body temperature, and (C) survival percentages are shown (n=4-5/group). Subsets of mice (n=3-6/group) were euthanized at 3 days post-infection (dpi); (D) lung viral titers were measured by TCID_50_ (dashed line, limit of detection), and (E-J) histopathology analysis was performed in the lungs by H & E staining. (E) Total inflammation scores and (F) parenchymal inflammation between the four virus-infected groups are presented. (G) Representative histopathology images of non-obese males (NO-M), obese males (O-M), non-obese females (NO-F), and obese females (O-F) are shown. Statistical comparisons were done using one-way ANOVA followed by Tukey’s post-hoc comparisons or Kruskal-Wallis test followed by Dunn’s post-hoc comparisons.

To test if the severe disease in females is associated with greater pulmonary inflammation, we performed histopathology analysis of H&E-stained lungs at 3 dpi. Inflammation in the pleura, lung airways, vessels, and parenchyma was evaluated. While there were no obvious lesions in lungs from medium-inoculated mice, bronchopneumonia and arteritis were evident in IAV-infected mouse lungs (**Supplementary Figures 4A-B**). Among virus-infected mice, there was no significant difference in total lung inflammation score (**Figure 2E**). Likewise, inflammatory responses were comparable among the four groups in the pleura, lung airways, and pulmonary vessels **(Supplementary Figures 4C-E)**. Interestingly, inflammation in the lung parenchyma was more severe in female mice with obesity compared to all other groups, as indicated by the average inflammation scores (i.e., 1.2 for NO-M, 2 for NO-F, 1.6 for O-M, and 3.3 for O-F, out of the maximum score of 5) **(Figures 2F-J)**. These data suggest that after high-dose infection, female mice, both non-obese and obese, exhibited shorter survival time than males, and it was not driven by lung virus titers, which were comparable across all groups. Females with obesity, however, exhibited more severe pulmonary parenchymal inflammation than other groups

### Pulmonary cytokine and chemokine induction at 3 dpi varied by biological sex and obesity status after a high-dose IAV infection

To further compare inflammatory changes, we measured 26 cytokines and chemokines in the lung homogenates at 3 dpi. Infection with high-dose IAV induced production of various cytokines and chemokines in the lungs of all groups, including GM-CSF, GRO-α, IL-6, TNF-α, IL-1β, IFN-γ, IL-18, MCP-1, MCP-3, MIP-1α, MIP-1β, MIP-2α, and IP-10, irrespective of biological sex and obesity status (**Supplementary Figure 5**). However, concentrations of IL-12p70, IL-27, and RANTES showed a higher trend or significant difference in virus-infected mice compared to medium-inoculated mice of the other three groups, except for non-obese males. Likewise, IL-22 showed a higher trend only in virus-infected males and females with obesity but not in non-obese mice **(Supplementary Figure 5)**. When the absolute concentrations of cytokines and chemokines were compared among virus-infected groups, a statistical difference was observed only for Eotaxin and IL-5, which were both significantly lower in virus-infected non-obese females compared to non-obese males and females with obesity (**Supplementary Figure 5**).

To understand how high-dose virus infection impacts relative changes in the inflammatory responses, we calculated the fold change in pulmonary cytokines and chemokines in male and female mice, with or without obesity, compared to their age- and sex-matched medium-inoculated controls. Sex difference was observed in IL-1β fold change, where females, irrespective of obesity status, had significantly higher induction compared to the males (**Figure 3A**). Likewise, males and females with obesity had a higher trend of IL-1β than their non-obese controls. Likewise, the effect of obesity was observed in IL-17A expression, where mice with obesity, irrespective of sex, had significantly higher expression compared to non-obese controls (**Figure 3B**). The male-specific effect of obesity was observed in TNFα and IL-6 induction, which was significantly higher and trended higher respectively in males with obesity compared to the non-obese males, but no such difference was observed in females (**Figure 3C-D**). IL-6 induction trended higher in non-obese females compared to non-obese males. IL-2 and IL-23-fold changes, however, were predominantly higher only in females with obesity compared to other groups (**Figure 3E-F**). Non-obese females had significantly higher IL-12p70 induction compared to non-obese males, with no such difference observed among mice with obesity **(Figure 3G)**. Sex difference was observed in IL-5 induction which was either significantly lower or trended low in females compared to males **(Figure 3H)**. Fold change in IL-22 was significantly lower in non-obese males compared to the other three groups, which had comparable induction (**Figure 3I**). In the analysis of the chemokines, MCP-1 fold change in non-obese females was significantly higher than non-obese males and both males and females with obesity; MIP-1α fold change was significantly higher in females than in males and mice with obesity had higher induction than non-obese mice; and the fold change of GRO-α showed a higher trend in females with obesity compared to males with obesity (**Figure 3J-L**). These data indicate unique sex-specific and obesity-specific alterations in pulmonary cytokines and chemokines after infection with a high-dose IAV.

**Figure 3.**
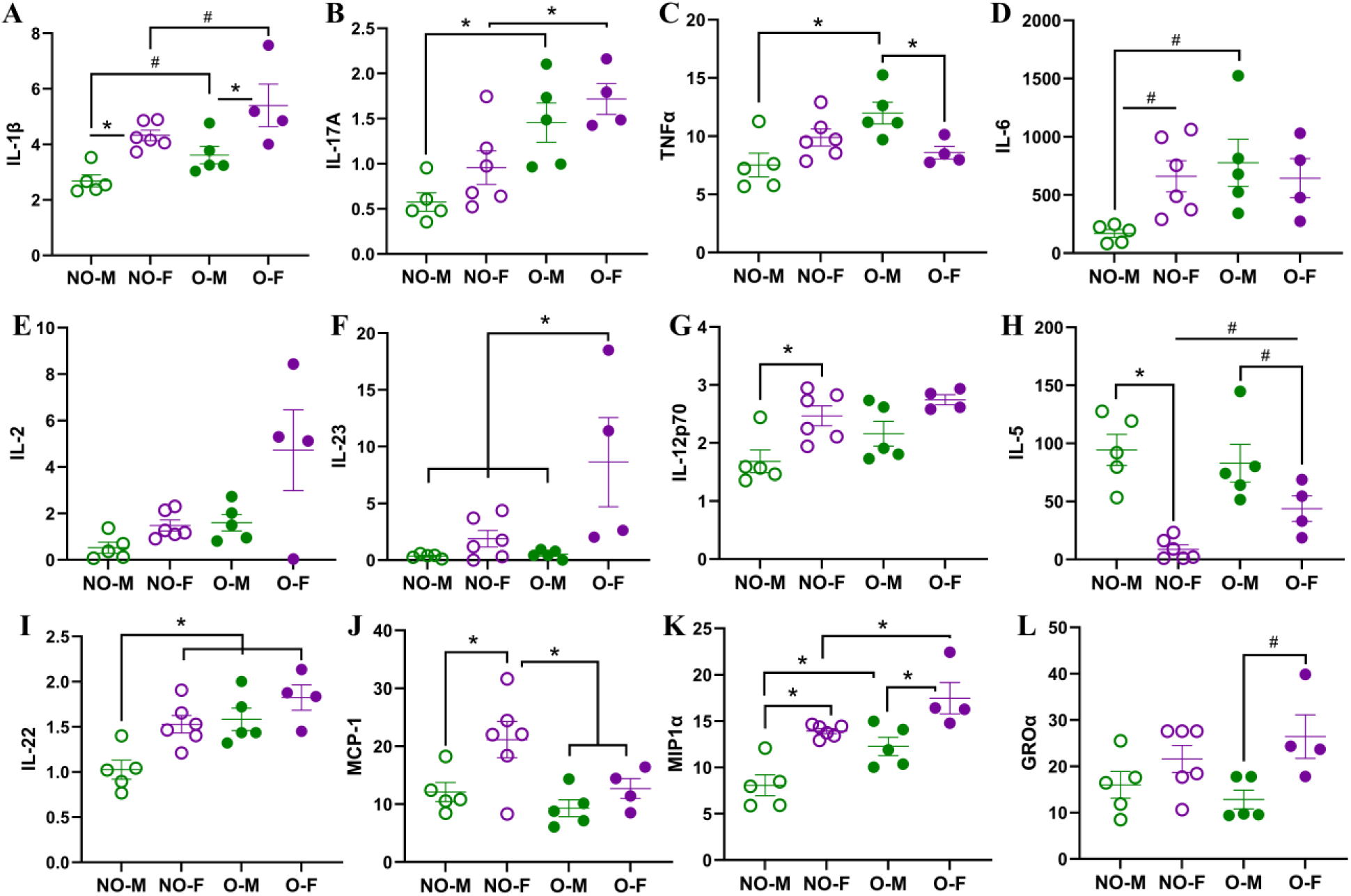
Cytokine and chemokine responses in the lungs following a high-dose IAV infection. After medium or high-dose IAV infection, mice were euthanized at 3 days post-infection (dpi), and cytokines and chemokines were measured in lung homogenates. Fold changes in cytokines and chemokines in virus-infected non-obese males (NO-M), non-obese females (NO-F), obese males (O-M), and obese females (O-F), compared to their medium-inoculated controls are shown for (A) IL-1β, (B) IL-17A, (C) TNFα, (D) IL-6, (E) IL-2, (F) IL-23, (G) IL-12p70, (H) IL-5, (I) IL-22, (J) MCP-1, (K) MIP1α, and (L) GROα. Data is shown as mean ± standard error of the mean (SEM) (n=3-6/group). Statistical comparisons were made using one-way ANOVA or Kruskal-Wallis test, and p-values were adjusted for multiple comparisons using the Benjamini-Hochberg method. An asterisk (*) indicates a statistically significant difference (p<0.05) while a hash (^#^) indicates a trend (0.05≤p≤0.1).

### After a low-dose influenza virus infection, females with obesity suffered from more severe disease, which was not associated with lung virus titers

After infection with a low dose of IAV (i.e., 10^1.5^ TCID_50_), the absolute (from 10-21 dpi) and relative (on 15, 17 to 21 dpi) body mass losses were significantly higher in males with obesity compared to non-obese males (**Figure 4A-B**). A similar effect of obesity was observed in females, where the absolute (from 13-21 dpi) and relative (from 14-21 dpi) body mass losses were significantly higher in females with obesity compared to non-obese females (**Figure 4A-B**). We also compared the biological sex differences in body mass loss. The percentage change in body mass was significantly greater in non-obese females from 9-13 dpi compared to non-obese males, while it was significantly greater in females with obesity compared to males with obesity between 14-15 dpi (**Figure 4B**). Though peak body mass loss occurred between 9-13 dpi, and females, regardless of obesity status, experienced the greatest losses, non-obese mice recovered to their baseline body mass by 21 dpi, whereas mice with obesity did not. The percentage change in body temperature also showed a similar trend where mice with obesity, irrespective of sex, lost significantly greater body temperature than the non-obese mice; and females, with or without obesity, lost significantly greater temperature, indicating more severe disease (**Figure 4C**). These data suggest that obesity is associated with increased disease severity following IAV infection in both males and females. Furthermore, females, irrespective of obesity status, suffer greater body mass loss after infection. Overall, females with obesity exhibited the most severe outcome, which is evident from the observation that even at a low-dose infection, 2 of 8 (i25%) females with obesity lost ≥25% of their original body mass, and were humanely euthanized (**Figure 4D**) while all mice from other groups survived. Overall, these data indicate that obesity results in severe morbidity after infection with a low dose of IAV, and the effect is more severe for females with obesity.

**Figure 4.**
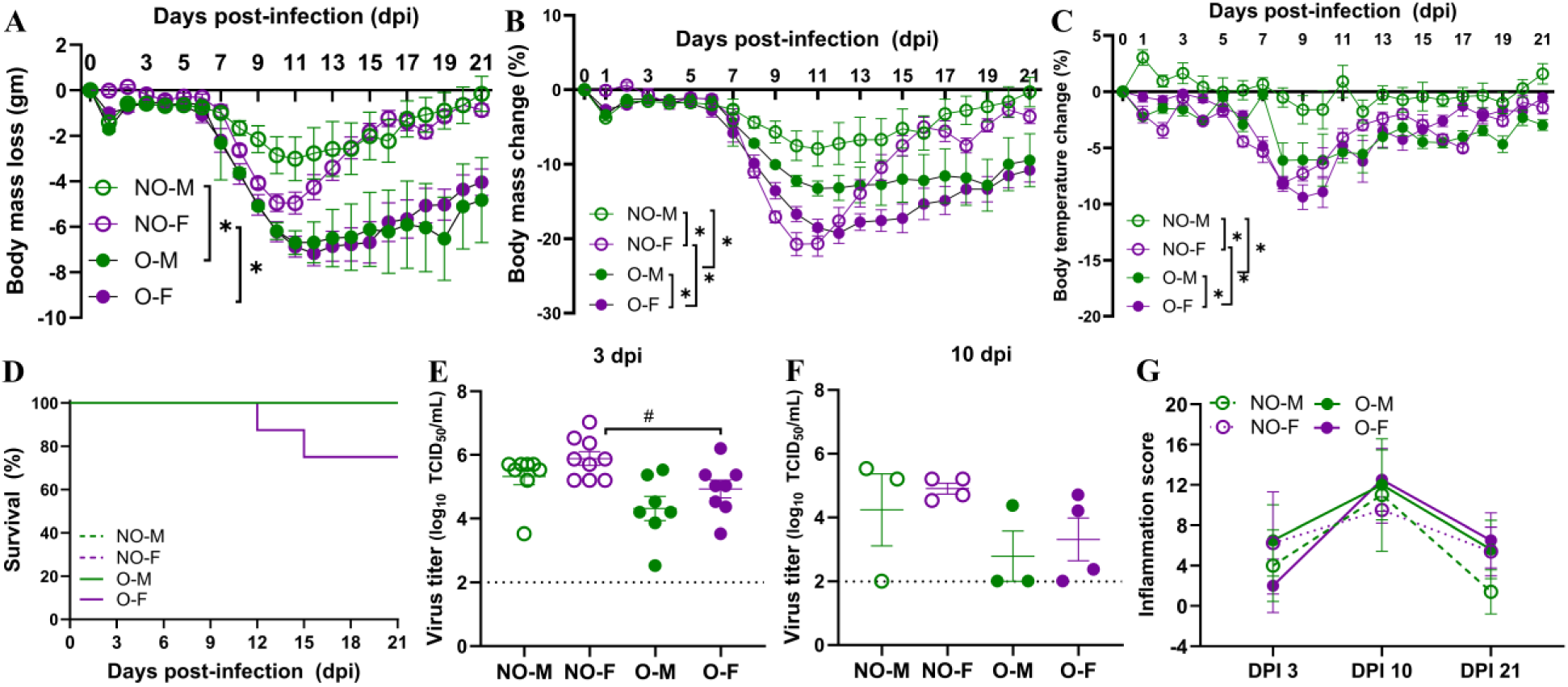
Morbidity and virus replication following a low-dose IAV infection. Male and female mice, with or without obesity, were intranasally inoculated with a low dose (i.e., 10^1.5^ TCID_50_) of the 2009 pandemic H1N1 IAV and monitored for 21 days post-infection (dpi) for changes in body mass and temperature. (A) Absolute (gm) and (B) relative (%) changes in body mass and (C) relative (%) change in body temperature, and (D) survival curves are compared (n=5-8/group). Virus titers in the lungs were measured by TCID_50_ assay at (E) 3 dpi (n=7-9/group) and (F) 10 dpi (n=3-4/group). Likewise, lung inflammation was assessed by hematoxylin and eosin (H&E) staining, and (G) total inflammation scores at 3, 10, and 21 dpi are compared. Data is shown as mean ± standard error of the mean (SEM). Statistical comparisons were done using one-way ANOVA followed by Tukey’s post-hoc comparisons or Kruskal-Wallis test followed by Dunn’s post-hoc comparisons. Repeated measurements in A-C were compared by repeated measures ANOVA (mixed model) and Tukey’s multiple comparisons. An asterisk (*) indicates a statistically significant difference (p<0.05) while a hash (^#^) indicates a trend (0.05≤p≤0.1). Abbreviations: NO-M, non-obese males; NO-F, non-obese females; O-M, obese males; and O-F, obese females.

To determine if the difference in viral replication in the lungs contributes to the differences in disease severity among different groups, we euthanized subsets of mice infected with a low dose of IAV at 3 dpi and collected lung tissue for virus titration. Interestingly, there was a lower trend (p=0.1) of virus titers in the lungs of females with obesity as compared to the non-obese females (**Figure 4E**). Virus was detected in the lungs of mice even at 10 dpi, with no statistical difference observed (**Figure 4F**). These data suggest that increased disease severity in mice with obesity and higher morbidity in females with obesity are not only driven by the virus replication in the lungs.

### Female mice with obesity had delayed but persistent histopathological changes after a low-dose IAV infection

After a low-dose IAV infection, mice were euthanized at 3, 10, and 21 dpi, and pulmonary pathological changes were compared by H&E staining. When total inflammation scores were compared among the groups, there were no significant differences at 3, 10, and 21 dpi. However, an interesting trend was observed where females with obesity displayed the lowest inflammation score at 3 dpi compared to other groups, but the inflammation score was highest of all groups highest at 10 dpi and remained highest even at 21 dpi **(Figure 4G)**. Non-obese females, who lost body mass comparable to females with obesity during the peak disease **(Figures 4A-B)**, had a greater inflammation score at 3 dpi that increased further at 10 dpi and decreased at 21 dpi **(Figure 4G)**.

Then, we compared the inflammatory changes within the same group over time (**Figure 5**). In non-obese males, inflammation was observed at 3 dpi (average score: 4), peak inflammation was at 10 dpi (average score: 11), and there was a significant decline in inflammation by 21 dpi (average score: 1.4) (**Figure 5A**). Representative images of non-obese males at 3 dpi showing inflammation mainly centered on vessels (V); 10 dpi showing inflammation centered on airways and arteries; and 21 dpi showing residual parenchymal inflammation around the main stem bronchus (MSB) are shown (**Figure 5A**). In non-obese females, inflammation was relatively stronger at 3 dpi (average score: 6.3); reached a peak at 10 dpi (average score: 9.5); and there was a lower trend by 21 dpi (average score: 5.4) (**Figure 5B**). Representative images of non-obese females at 3 and 10 dpi, showing multifocal inflammation, and at 21 dpi with residual inflammation are shown (**Figure 5B**). In males with obesity, inflammation was relatively stronger at 3 dpi (average score: 6.5), peaked at 10 dpi (average score: 12), and there was a lower trend by 21 dpi (average score: 5.6) (**Figure 5C**). Representative images of males with obesity at 3, 10, and 21 dpi are shown (**Figure 5C**). Interestingly, in females with obesity, inflammation at 3 dpi was relatively weaker (average score: 2); significantly increased at 10 dpi (average score: 12.5); and there was a significant reduction in 21 dpi compared to 10 dpi but inflammation at 21 dpi was still having a higher trend than 3 dpi (average score: 6.5) (**Figure 5D**). Representative images for females with obesity at 3, 10, and 21 dpi are shown (**Figure 5D**). Overall, these data suggest that groups other than non-obese males had residual inflammation even at 21 dpi. Females with obesity had a delayed onset of pulmonary pathological changes that increased significantly at 10 dpi and persisted longer up to 21 dpi, correlating with severe morbidity and delayed recovery.

**Figure 5.**
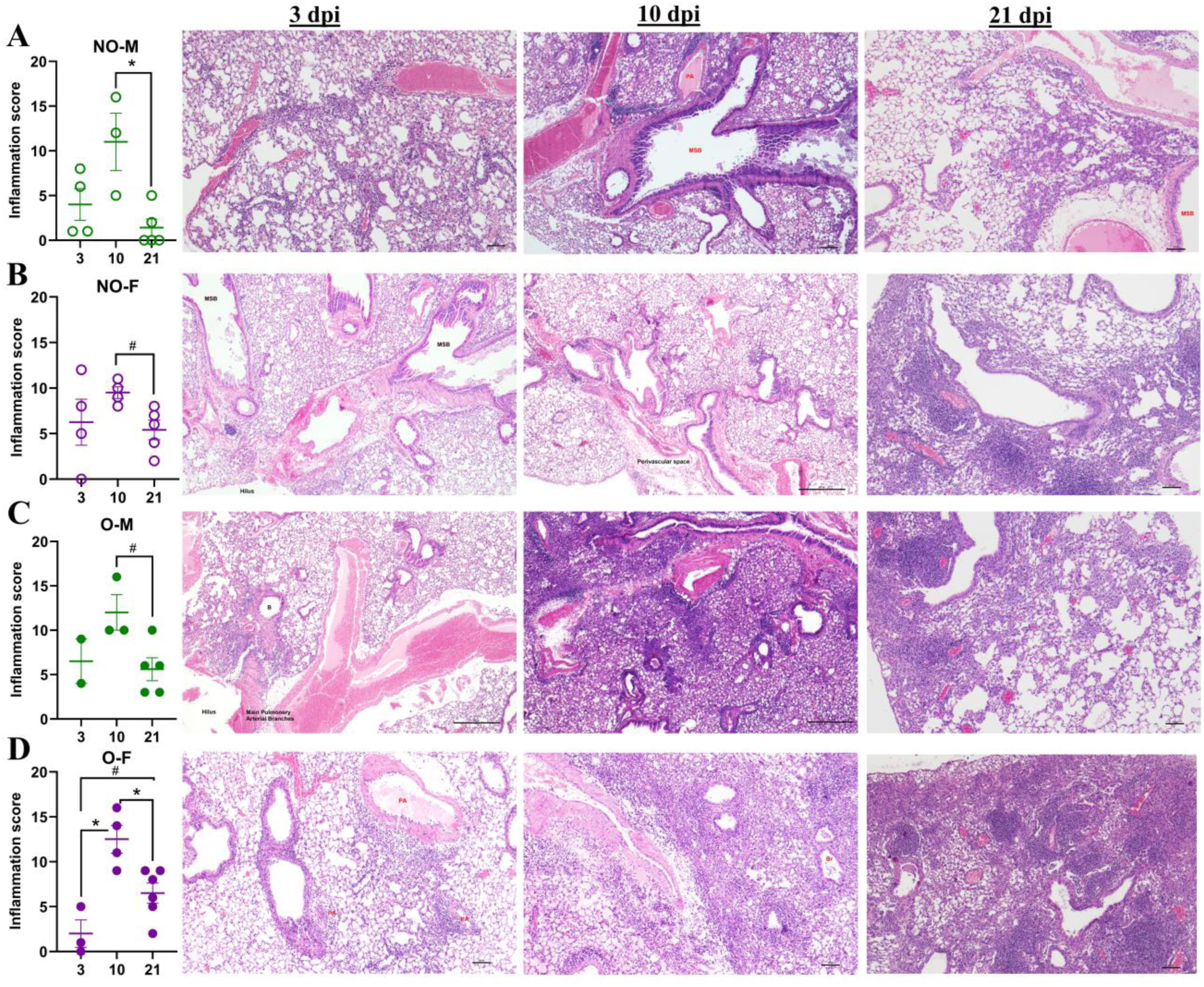
Progression of lung inflammation following low-dose IAV infection. Male and female mice, with or without obesity, were intranasally inoculated with a low dose (i.e., 10^1.5^ TCID_50_) of the 2009 pandemic H1N1 IAV. Subsets of mice were euthanized at 3, 10, and 21 dpi, and histopathology analysis by hematoxylin and eosin (H&E) staining was performed. Total inflammation scores at 3, 10, and 21 dpi together with their representative images for the (A) non-obese males (NO-M), (B) non-obese females (NO-F), (C) males with obesity (M-O), and (D) females with obesity (F-O) are shown (n=2-6/group). Data is shown as mean ± standard error of the mean (SEM). Statistical comparisons were performed using the one-way ANOVA or Kruskal-Wallis test followed by Tukey’s or Dunn’s multiple comparisons, respectively. An asterisk (*) indicates a statistically significant difference (p<0.05) while a hash (^#^) indicates a trend (0.05≤p≤0.1). Abbreviations – V: vessel, PA: pulmonary artery, MSB: main-stem bronchi, and Br: bronchiole.

### Female mice with obesity suffered from delayed but persistent heightened pulmonary cytokine and chemokine influx following a low-dose IAV infection

After the low-dose virus infection, we measured 48 cytokines and chemokines in the lung homogenates at 3 dpi, 10 dpi, and 21 dpi. At 3 dpi, the absolute concentrations of cytokines and chemokines were higher in non-obese females than in the other groups (**Supplementary Figure 6A**). Non-obese females had significantly higher or higher trend of IL-28 (known as IFN-λ), IL-18, IL-2, IL-2R, IL-4, IL-7, IL-7Rα, IL-9, TNFα, IL-10, IL-19, IL-12p70, IL-27, IL-17A, LIF, MCP-1, MIP-1α, MIP-1β, RANTES, BTC, GM-CSF, G-CSF, IL-5, and IL-13 compared to non-obese males. Likewise, non-obese females also had either significantly higher or a higher trend of IL-28, IL-18, IL-2, IL-2R, IL-4, IL-7, IL-7Rα, IL-9, TNFα, IL-10, IL-12p70, IL-27, IL-17A, LIF, MIP-1α, MIP-1β, BTC, GM-CSF, and IL-13 compared to females with obesity. Though the concentrations of IL-6 and IL-31 were higher in non-obese females than in the other groups, they were not significant (**Supplementary Figure 6A**).

At 10 dpi, IL-1α and IL-33 were significantly lower in males with obesity compared to non-obese males (**Supplementary Figure 6B**). IL-1α and RANKL were also significantly lower in males with obesity compared to females with obesity. Non-obese females, which had elevated levels of cytokines/chemokines at 3 dpi, maintained these concentrations, while obese females, which had a lower concentration of cytokines/chemokines at 3 dpi, had them elevated at 10 dpi (**Supplementary Figure 6A-B**). Females, with or without obesity, had comparable cytokines and chemokines at 10 dpi except for significantly higher concentration of IL-1α observed in females with obesity compared to the non-obese females (**Supplementary Figure 6B**).

At 21 dpi, males and females with obesity maintained higher inflammatory cytokines and chemokines. The concentrations of IL-2 and IL-31 were significantly lower and IL-3 and IL-17A trended lower in males with obesity compared to non-obese males (**Supplementary Figure 6C**). The concentration of GRO-α was significantly higher in males and females with obesity compared to non-obese mice, and a higher trend was observed in non-obese males than non-obese females. Females with obesity had significantly higher concentrations of MCP-1 and IFN-γ but a lower trend of MIP-2α compared to non-obese females. The concentration of IP-10 also showed a higher trend in females with obesity compared to males with obesity and non-obese females. Females with obesity also had a significantly higher IL-2 and a higher trend of IL-13 compared to males with obesity. At 3, 10, and 21 dpi, leptin concentrations were significantly higher in males and females with obesity compared to non-obese controls (**Supplementary Figure 6C**).

The comparison of fold changes further illustrated the differences in the cytokines and chemokines more clearly (**Figure 6A-C**). At 3 dpi, non-obese females had either significantly higher fold changes or higher trend for IL-28, IL-1α, IL-18, IL-1β, IL-2, IL-2R, IL-4, IL-7, IL-7Rα, IL-9, IL-15, TNFα, RANKL, BAFF, IL-10, IL-19, IL-22, IL-12p70, IL-23, IL-27, LIF, MCP-1, MIP-1α, MIP-1β, RANTES, MCP3, eotaxin, IP-10, G-CSF, GM-CSF, M-CSF, VEGF-A, and IL-13 compared to non-obese males; and that of IL-28, IL-1α, IL-18, IL-2, IL-2R, IL-4, IL-7Rα, IL-15, TNFα, RANKL, BAFF, IL-10, IL-27, LIF, MCP-1, MIP-1α, MIP-1β, MCP-3, eotaxin, IP-10, BTC, G-CSF, GM-CSF, and M-CSF compared to females with obesity. Females with obesity had higher trend or significant fold changes of IL-18, IL-33R, IL-2, IL-4, IL-7, IL-7Rα, IL-9, IL-10, IL-19, IL-23, IL-27, LIF, BTC, and VEGF-A compared to males with obesity. Females with obesity also had significantly higher induction of BTC than non-obese females. Males with obesity had significantly higher fold changes of IL-28 and RANTES; and higher trend for IL-2 and IL-13 compared to non-obese males, while Eotaxin had higher trend in non-obese males (**Figure 6A**). At 10 dpi, non-obese females had significantly higher induction of IL-1α, IL-7, IL-9 and a higher trend for MIP-1α and eotaxin compared to non-obese males, and significantly higher MCP-3 than females with obesity (**Figure 6B**). Females with obesity had significantly higher induction of IL-1α, IL-7, IL-7Rα, IL-9, RANKL, MIP-1α and M-CSF and higher trend for IL-23, IL-27, BTC, GM-CSF but a significantly lower IL-33 induction compared to males with obesity. Females with obesity also had significantly higher induction of IL-7, IL-9, MCP-3, and a higher trend of IL-1α, IL-13, and BTC compared to the non-obese females. Non-obese males had a higher trend of IL-1α and IL-2 compared to males with obesity, while males with obesity had significantly higher induction of IL-33 and RANTES and higher trend for eotaxin than in non-obese males (**Figure 6B**).

**Figure 6.**
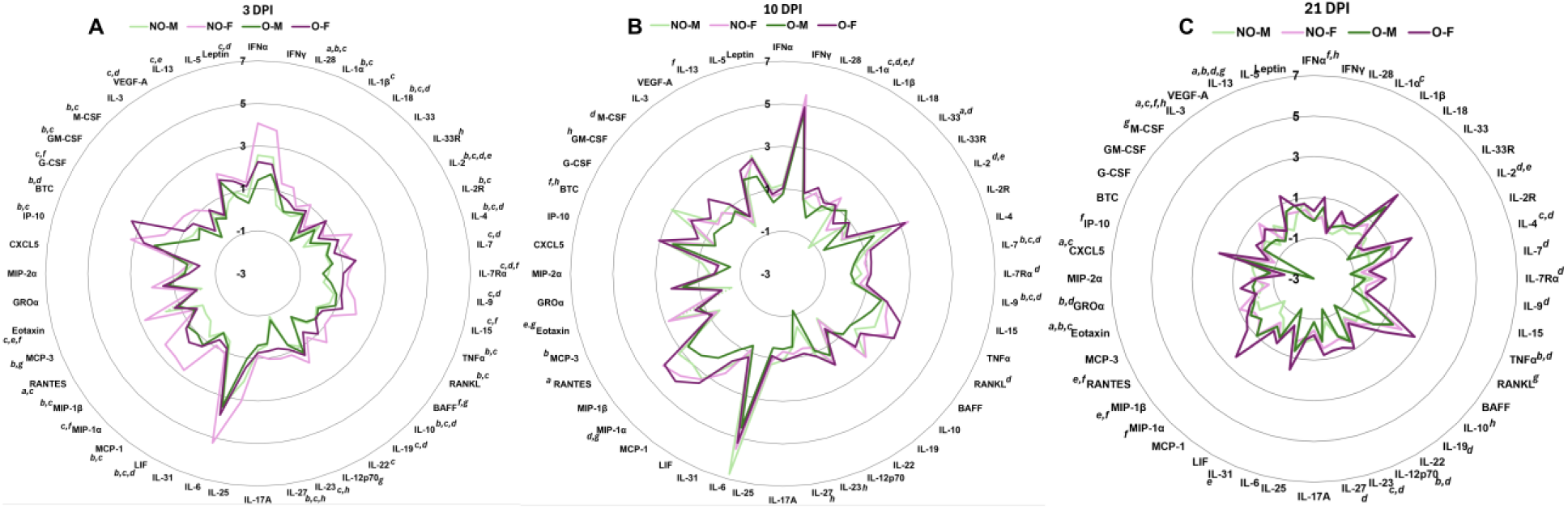
Cytokine/chemokine responses in the lungs following low-dose IAV infection. After infection with a low dose (i.e., 10^1.5^ TCID_50_) of the 2009 pandemic H1N1 IAV, subsets of mice were euthanized at 3-, 10-, and 21-days post-infection (dpi), lungs were collected, and various cytokines and chemokines were measured. Fold changes in cytokines and chemokines in virus-infected mice compared to their medium-inoculated controls are compared. Data for (A) 3 dpi, (B) 10 dpi, and (C) 21 dpi are shown. Statistical comparisons were made using one-way ANOVA or Kruskal-Wallis test, and p-values were adjusted for multiple comparisons using the Benjamini-Hochberg method. In figures, a and e; b and f; c and g; and d and h represented a significant difference or a trend between non-obese males (NO-M) versus obese males (O-M); non-obese females (NO-F) versus obese females (O-F); non-obese males versus non-obese females; and obese males versus obese females, respectively.

At 21 dpi, females with obesity maintained a higher induction of cytokines/chemokines than the other groups (**Figure 6C**). Fold changes in IFN-α, IL-2, IL-4, IL-7, IL-7Rα, IL-9, TNF-α, IL-10, IL-12, IL-19, IL-23, IL-27, GRO-α, IL-3, and IL-13 were either significantly higher or had a higher trend in females with obesity compared to males with obesity. Females with obesity also had significantly greater induction or higher trend of IFNα, TNFα, IL-12p70, MIP-1α, MIP-1β, RANTES, GROα, IP-10, IL-3, and IL-13 compared to non-obese females. Non-obese females had significantly higher induction or higher trend for IL-1α, IL-4, IL-23, RANKL, eotaxin, M-CSF, and IL-13 than non-obese males and a significantly higher induction of eotaxin than females with obesity. Non-obese males had a higher trend or significant induction of IL-2, IL-31, CXCL-5, and IL-3 than males with obesity, and a significantly higher IL-3 and CXCL-5 than non-obese females. Males with obesity, on the other hand, had significantly higher induction of eotaxin and IL-13 and a higher trend of MIP-1β and RANTES compared to non-obese males. These data suggest that in non-obese females, cytokines and chemokines were induced at high levels at 3 dpi, which was maintained up to 10 dpi, and declined by 21 dpi; while in females with obesity, the levels were lower at 3 dpi, reached comparable levels to non-obese females at 10 dpi, but remained higher even at 21 dpi. The cytokine and chemokine data corroborate with the histopathology data, indicating delayed but persistent inflammation in females with obesity.

### Pulmonary myeloid and lymphoid cell frequencies during the acute phase are lower in females with obesity

At 3 dpi, frequencies of myeloid and lymphoid cells were determined in the lungs. CD45^+^ cells had a lower trend in non-obese males compared to non-obese females and males with obesity, and there was no difference in the frequencies of neutrophils and eosinophils (**Figure 7A-C**). The frequencies of alveolar macrophages, M1 and M2 macrophages, and Ly6c^+^ inflammatory monocytes were all higher in non-obese females compared to other groups but did not reach statistical significance (**Figure 7D-J**). The interstitial macrophages, including the M1 types, showed a higher trend in non-obese females than in females with obesity (**Figure 7K-M**). There was no difference in CD11b DC frequencies, while the frequencies of moDCs were significantly higher in non-obese females compared to the females with obesity and showed a higher trend than in non-obese males (**Figure 7N-O**).

**Figure 7.**
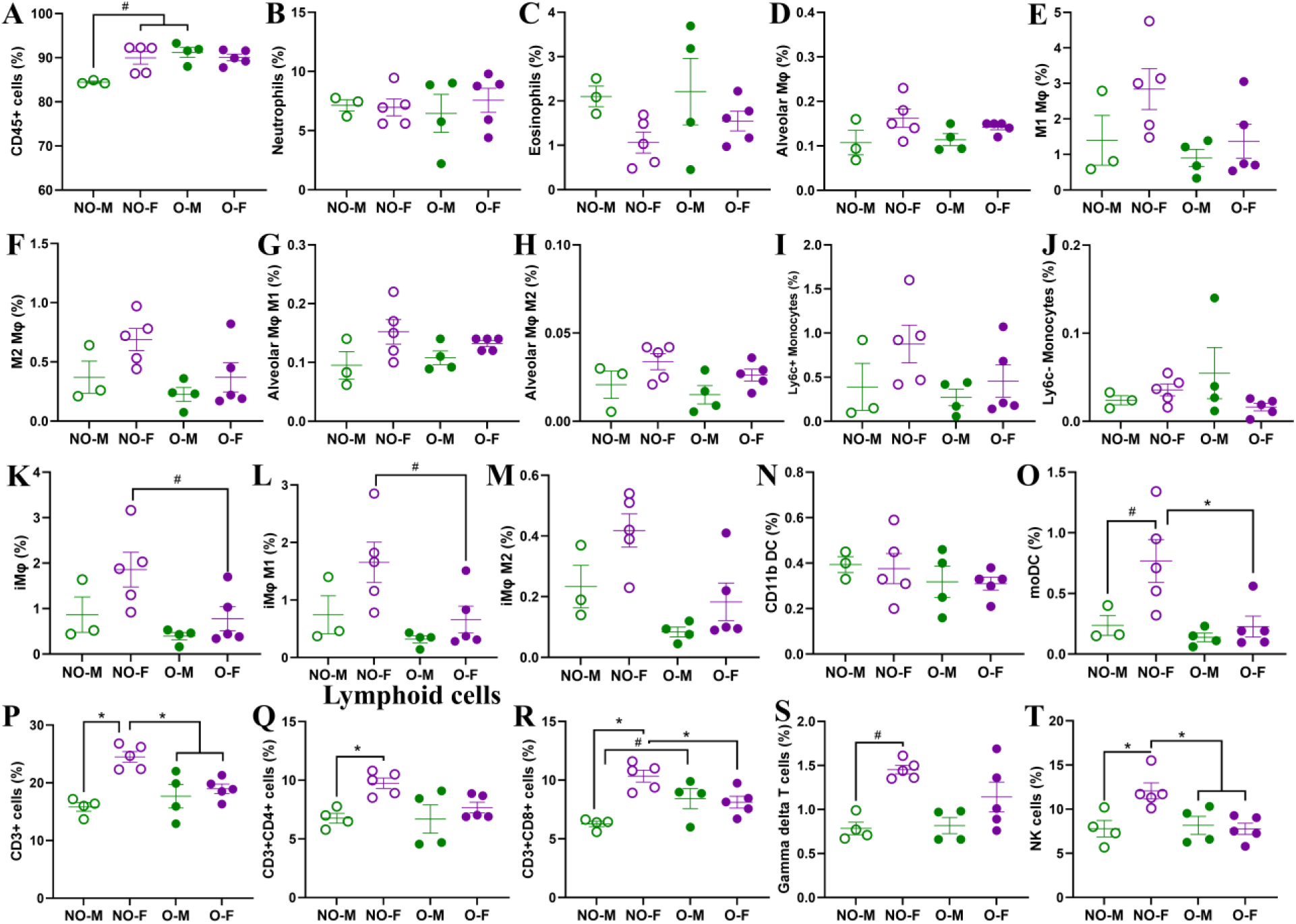
Characterization of myeloid and lymphoid cells in the lungs following a low-dose IAV infection. After infection with a low dose (i.e., 10^1.5^ TCID_50_) of the 2009 pandemic H1N1 IAV, subsets of mice were euthanized at 3 days post-infection (dpi) to determine the frequencies of various myeloid and lymphoid cells in the lungs. In myeloid panel, the frequencies of (A) CD45^+^ cells, (B) neutrophils, (C) eosinophils, (D) alveolar macrophages, (E) M1 macrophages, (F) M2 macrophages, (G) M1 and (H) M2 type alveolar macrophages, (I) Ly6c^+^ and (J) Ly6c^-^ monocytes, (K) interstitial macrophages, (L) M1 and (M) M2 type interstitial macrophages, (N) CD11b dendritic cells, and (O) monocyte-derived dendritic cells are compared. Likewise, frequencies of (P) CD3^+^ T cells, (Q) CD3^+^CD4^+^ T helper cells, (R) CD3^+^CD8^+^ T cells, (S) gamma delta T cells, and (T) NK cells are compared. Data is shown as mean ± standard error of the mean (SEM) (n=3-5/group). Statistical comparisons were performed using the one-way ANOVA or Kruskal-Wallis test, followed by Tukey’s or Dunn’s multiple comparisons, respectively. An asterisk (*) indicates a statistically significant difference (p<0.05) while a hash (^#^) indicates a trend (0.05≤p≤0.1). Abbreviations: NO-M, non-obese males; NO-F, non-obese females; O-M, obese males; and O-F, obese females.

Among the lymphoid cells, the frequencies of CD3^+^ cells were significantly higher in non-obese females compared to the other groups (**Figure 7P**). The frequency of T helper (CD3^+^CD4^+^) cells was also significantly higher in non-obese females compared to non-obese males (**Figure 7Q**). The CD3^+^CD8^+^ T cells were significantly higher in non-obese females compared to non-obese males and females with obesityMales with obesity also had a higher trend of CD3+CD8+ T cells compared to non-obese males (**Figure 7R**). There was no difference in the frequencies of regulatory T cells at 3 dpi in the lungs. However, the gamma delta T cells were highest in the non-obese females, which showed a higher trend compared to non-obese males. Likewise, the NK cell frequency was significantly higher in non-obese females compared to all other groups (**Figure 7S-T**). Together, these data suggest that deposition of myeloid and lymphoid cells was greater in non-obese females compared to the other groups, including the females with obesity.

### Females with obesity induced antibodies equivalent to non-obese females and higher than males with obesity at 21 dpi following low-dose IAV infection

At 21 dpi, systemic and mucosal antibody responses were compared in plasma (**Figure 8A-D**) and lung homogenates (**Figure 8F-H**). Sex difference was observed in systemic antibody responses, where females had higher antibodies than males **(Figures 8A-D)**. In plasma, IgG and IgG2c antibodies were significantly higher in non-obese females compared to non-obese males, and females with obesity had significantly higher IgG and a higher trend of neutralizing antibodies (nAb) than males with obesity. In the lungs, B-cell frequency was determined by flow cytometry at 3 dpi. B-cell frequency was significantly lower in the lungs of non-obese female mice at 3 dpi compared to non-obese males and females with obesity (**Figure 8E**). However, females, irrespective of obesity status, had higher local antibody responses in the lungs. IgG antibody in non-obese females was significantly higher than in non-obese males, while the IgA and nAb titers were also higher but not significant (**Figure 8F-H**). Females with obesity also produced higher IgG, IgA, and nAb titers in the lungs compared to the males with obesity. While 100% of females with obesity had higher neutralizing antibodies in the lungs, 40% of males with obesity had no detectable neutralizing antibodies **(Figures 8F-H).** Overall, a female-biased higher antibody induction was observed following primary influenza virus infection, in both non-obese mice and mice with obesity, as evident from both local (lung) and systemic (plasma) samples.

**Figure 8:**
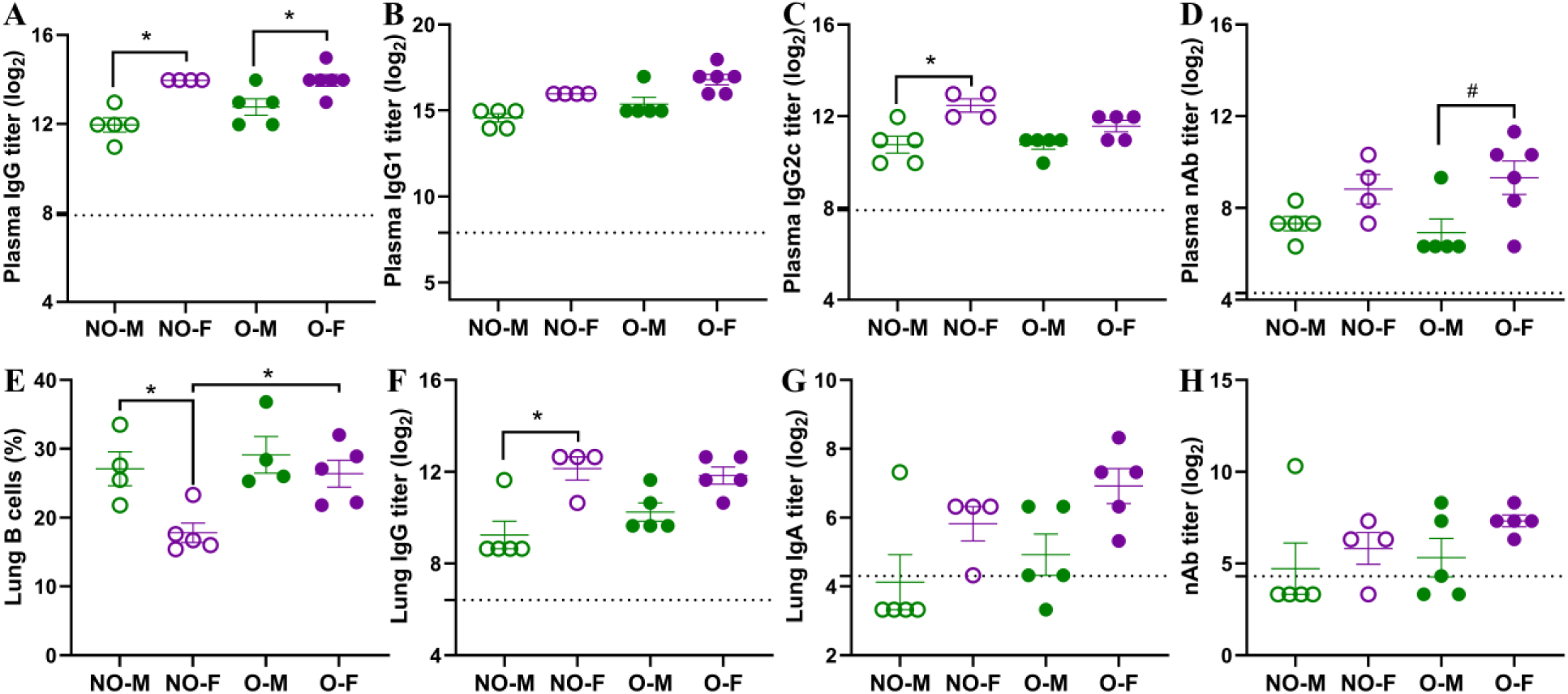
Antibody responses in plasma and lung homogenates following low-dose IAV infection. After infection with a low dose (i.e., 10^1.5^ TCID_50_) of the 2009 pandemic H1N1 IAV, mice were followed for morbidity measurements up to 21 dpi. At 21 dpi, after euthanization, plasma samples and lung homogenates were prepared to measure systemic and mucosal antibody responses. Plasma levels of (A) IgG, (B) IgG1, (C) IgG2c, and (D) virus-neutralizing antibody (nAb) titers are compared. (E) The frequency of B-cells in the lungs, as measured by flow cytometry in the subsets of mice euthanized at 3 dpi is shown. Likewise, pulmonary (F) IgG, (G) IgA, and (H) nAb titers are compared. Data is shown as mean ± standard error of the mean (SEM) (n=4-6/group). Statistical comparisons were made using one-way ANOVA or Kruskal-Wallis test followed by Tukey’s or Dunn’s multiple comparisons, respectively. An asterisk (*) indicates a statistically significant difference (p<0.05) while a hash (^#^) indicates a trend (0.05≤p≤0.1). Abbreviations: NO-M, non-obese males; NO-F, non-obese females; O-M, obese males; and O-F, obese females.

### Non-responder females on HFD had morbidity, virus titers, and pulmonary inflammation comparable to females with obesity

As some of the females on HFD were non-responders, we wanted to compare whether the outcomes of influenza virus infection would be similar between the non-responder and responder (i.e., obese) females. The body mass, BMI, and GTT values were significantly lower in non-responder females compared to the females with obesity (**Figure 9A-C**). After a high-dose infection, the median survival time was 9 dpi for both groups (**Figure 9D**). After a low-dose infection, 20% of the obese females and 80% of the non-responders were euthanized as they reached the humane endpoint (**Figure 9E**). We also compared virus replication in the lungs after infection with the low dose at 3 dpi, and it was similar between the groups (**Figure 9F**). To determine if the induction of inflammatory cytokines and chemokines differs between the non-responder and responder groups, we also compared the 48 cytokines/chemokines at 3 dpi, and there were similar responses (**Figure 9G**). Only difference observed was in leptin concentration, which showed a higher trend in the females with obesity than in non-responder females. This data suggests that, despite not gaining body mass to define them as having obesity in this model, long-duration intake of HFD increased the risk of severe outcomes of IAV infection, comparable to females that became obese.

**Figure 9.**
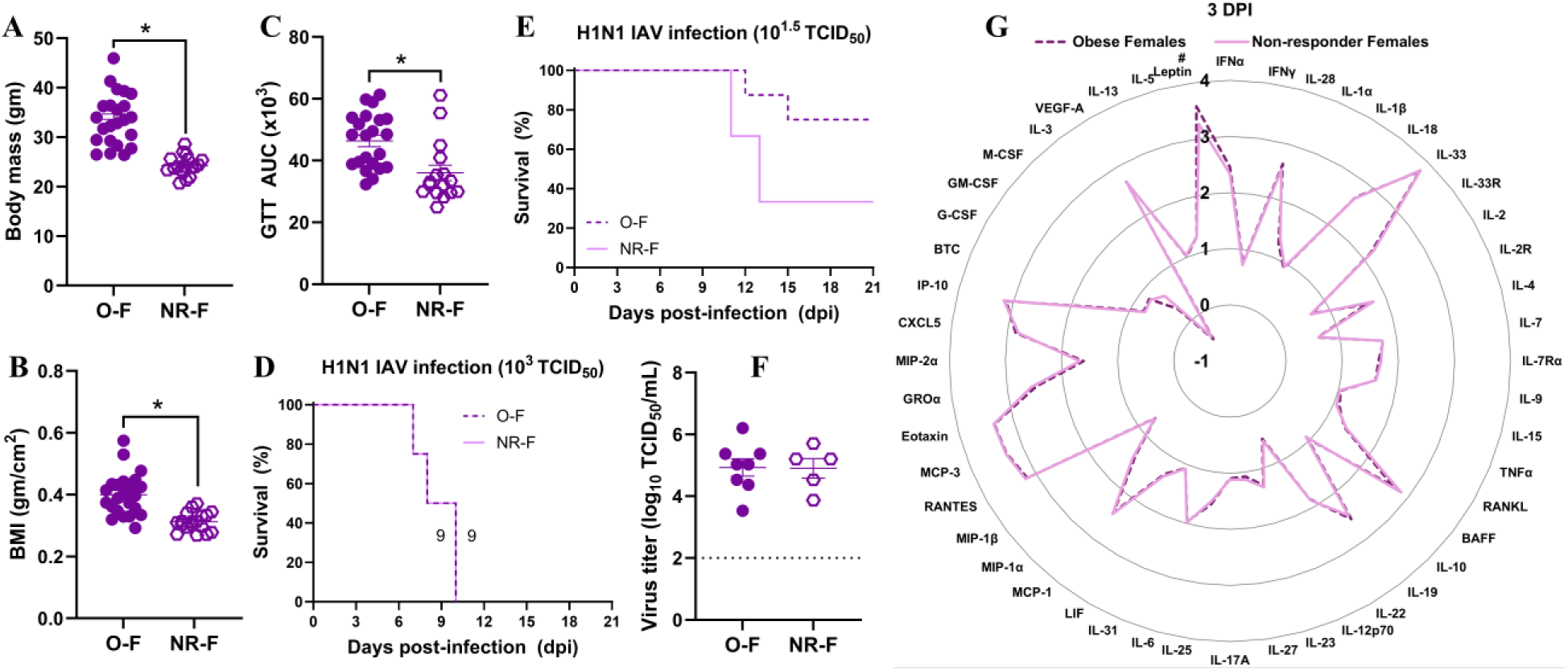
Comparison of morbidity, virus replication, and pulmonary cytokine and chemokine responses between HFD-treated responder and non-responder females. Female mice were on HFD for 13 to 14 weeks. About 30% of them were non-responders. (A) Body mass, (B) body mass index (BMI), and (C) glucose tolerance test (GTT) area under the curve (AUC) between females with obesity (i.e., the responders) and non-responder females are compared (n=16-23/group). Likewise, (D) survival rates after high-dose IAV infection and (E) low-dose IAV infection are shown (n=4-8/group). Similarly, (F) virus titers and (G) log_10_-transformed cytokine and chemokine concentrations in the lungs at 3 dpi with a low-dose virus infection are compared (n=5-8/group). Data is shown as mean ± standard error of the mean (SEM). Statistical comparisons were made using the unpaired t-test or Mann-Whitney test, and an asterisk (*) indicates a statistically significant difference (p<0.05) while a hash (^#^) indicates a trend (0.05≤p≤0.1).

## Discussion

In this study, we aimed to determine sex differences in influenza pathogenesis and disease severity during obesity using a DIO mouse model and both high- and low-dose infection models of the 2009 pandemic H1N1 IAV. Our findings suggest that sex difference is observed following influenza virus infection during obesity. Females with obesity suffer from more severe disease compared to males, and it is likely associated with the delayed but persistent pathological changes in the lungs.

After the HFD treatment, male mice developed obesity earlier than females. This is consistent with prior findings suggesting that female mice are more resistant to developing DIO compared to males (28, 29). Such resistance in females is likely associated with greater energy expenditure, estrogen-mediated protection, and compensatory increase in genes related to appetite control and energy balance in the hypothalamus (28, 30, 31). Female mice had a higher concentration of baseline and 14^th^ week adiponectin than the males. This might also contribute to slower body weight gain in females, as the adiponectin to leptin ratio is negatively correlated with body weight gain, adipose tissue deposition, and insulin resistance (32). In this study, we defined obesity as having a ≥20% body mass gain compared to the average body mass of age- and sex-matched mice on LFD treatment. Using this approach, around 70% of the females on HFD gained sufficient body mass to define them as having obesity. Females with obesity had greater body mass, BMI, adipose tissue deposition, glucose intolerance, and leptin concentrations compared to non-obese females; the observations were similar between males with or without obesity, making this approach a relevant model to study sex differences in IAV pathogenesis during obesity in both sexes.

Among non-obese mice, females exhibited greater influenza virus-driven disease severity. During the high-dose infection, female mice had a shorter median survival time (i.e., 8 vs 12 days), greater parenchymal inflammation, and higher induction of inflammatory cytokines and chemokines, including IL-1β, IL-6, IL-12p70, MCP-1, and MIP-1α, compared with males. Likewise, after a low-dose IAV infection, non-obese females exhibited significantly greater body mass loss and reduced body temperature, indicating higher disease severity than males. These findings are similar to earlier studies, which have shown that morbidity and mortality from H1N1 influenza virus infection are greater in lean females than in lean males (16, 22, 23, 33). Non-obese females had greater induction of several inflammatory cytokines and chemokines at 3 days post-infection, including IL-1β, TNFα, IL-12p70, MCP-1, MIP-1α, MIP-1β, IP-10, G-CSF, and GM-CSF associated with driving inflammation, lung pathology, and recruitment of neutrophils and monocytes in the lungs. This was also supported by the relatively higher deposition of myeloid and lymphoid cells in the lung tissues of non-obese females compared to non-obese males. The lung virus titers after both the high- and low-dose infection, however, were comparable between the non-obese males and females, indicating that observed sex differences in morbidity and mortality are not associated entirely with higher lung virus replication. Prior studies also suggest that sex differences during IAV infection are dose-dependent and although lung virus titers are comparable between males and females, the higher disease severity observed in non-obese females is associated with heightened inflammatory responses and pulmonary tissue damage (23, 34). Higher testosterone levels and amphiregulin, involved in lung tissue repair, are also associated with better influenza outcomes in non-obese males compared to non-obese females (23).

Although studies investigating sex differences during influenza and other viral infections among hosts with obesity are limited, Lee et al. observed that following infection with the alpha variant of SARS-CoV-2 in the K18-hACE2 mouse model of DIO, female mice with obesity exhibited more severe symptoms and higher inflammatory lung burdens than males (35). Prior studies of influenza pathogenesis in mouse models of DIO used either male-only (36–42), female only (43, 44), or males and females, but without stratification of data by sex (45, 46). One study examined both male and female DIO mice infected with a lethal dose of the virus and followed only up to 5 dpi, where they observed similar pathological changes after IAV infection in obese males and females (27). In our study, we investigated sex-specific phenotypes in obesity-associated influenza pathogenesis across different viral titers and over an extended period using both high- and low-dose infection models and following mice up to 21 dpi.

In our study, after infection with the high-dose IAV, females with obesity had a shorter median survival time (9 versus 13 days) than males with obesity. Despite having comparable lung virus titers and total pulmonary inflammation, the parenchymal inflammation was more severe in females than in males with obesity. Female mice also exhibited a stronger proinflammatory cytokine and chemokine response, including robust induction of IL-1β, MIP-1α, and GRO-α, suggesting greater recruitment of immune cells. Females with obesity also had robust induction of IL-23 and IL-17A cytokines, compared to males with obesity. These are consistent with the concept that activation of IL-23/IL-17 axis can contribute to heightened tissue inflammation and morbidity; however, definitive testing of this hypothesis will be required (47, 48).

Hornung et al. demonstrated that mice with obesity had a delayed onset of body mass loss compared to non-obese mice, and they did not recover to the baseline body mass until 21 dpi (43). Likewise, other studies have shown delayed mononuclear cell infiltration in the lungs, altered T cell responses, and a late rise in cytokine and chemokine responses in mice with obesity (49, 50). In our study, both males and females with obesity lost significantly greater body mass compared to their non-obese controls and did not recover to the baseline body mass even at 21 dpi, indicating the obesity-associated disease severity in both sexes. This reflects sustained inflammatory response and immune activation in mice with obesity, which may be responsible for the lack of weight recovery in mice with obesity. When comparing males and females with obesity, females lost greater body mass and temperature than males, and 25% of the females even reached the humane endpoint. The sex difference in low-dose IAV infection during obesity was not directly reflected by differences in lung virus replication. In both sexes, lung pathology peaked at 10 dpi and decreased at 21 dpi, but the residual lung inflammation was relatively higher in females with obesity. Females with obesity maintained robust induction of innate and inflammatory cytokines, including IFN-α, TNF-α, and GRO-α, even at 21 dpi when inflammation typically resolves during IAV infection. Likewise, compared to the males, females with obesity, maintained elevated levels of common gamma chain cytokines (i.e., IL-2, IL-4, IL-7, and IL-9), which sustain T- and B-cell activation and survival (51); IL-10 family cytokines (i.e., IL-10 and IL-19) with anti-inflammatory function; and IL-12 family cytokines (i.e., IL-12p70, IL-23, and IL-27) that activates Th1 and Th17 immune pathways (52). These observations suggest that females with obesity exhibit persistent inflammation even at 21 dpi, with inefficient inflammation resolution, indicating a pronounced dysregulated inflammatory response even compared to males with obesity (53). As in high-dose infection, persistent elevation of IL-23 during low-dose infection in females with obesity suggests a probable role for the IL-23/IL-17 inflammatory pathway in mediating severe influenza disease, warranting further exploration by direct mechanistic studies involving blockade by antibodies or genetic ablation (47, 48).

Females, irrespective of obesity, suffered from more severe disease following IAV infection compared to males. Following low-dose infection, females with or without obesity suffered similar levels of body mass loss at the peak disease. However, while non-obese females recovered from the severe disease, females with obesity could not. Interestingly, virus titer trended higher in non-obese females than the females with obesity, further suggesting the differences in disease severity to be mediated by host responses rather than lung virus titers. It is better explained by the delayed but persistent dysregulated inflammatory response in females with obesity than in non-obese females, as evidenced by lower frequencies of myeloid and lymphoid immune cell infiltration and lower pathological changes in the lungs at 3 dpi, and a switch from lower cytokines and chemokines induction at 3 dpi to higher induction at 21 dpi. In accordance, Smith et al. also showed reduced interstitial macrophages and dendritic cells in IAV-infected mice with obesity as compared to non-obese mice at 3 dpi (49). Reduced number and function of dendritic cells is also observed in humans with obesity (54). The non-responder females, treated with HFD but did not gain body mass to be defined as having obesity, also had disease severity, lung virus titers, and inflammatory changes comparable to females with obesity. These findings suggest that the metabolic effect of long-term HFD treatment may drive risk independently of overt obesity, and future studies should compare metabolic phenotypes before and after IAV infection between non-responder and responder females.

Following recovery from IAV infection, antibody responses are shown to be higher in non-obese females than in non-obese males, both in serum as well as in bronchoalveolar lavage (BAL) fluid (55). It is associated with a greater number of antibody-secreting B cells, T-helper cells, and germinal center B cells in the lungs of females than in males (55). We also observed greater systemic and mucosal antibody responses in non-obese females compared to non-obese males. Further, this study suggests that female sex drives higher antibody responses following recovery from IAV infection, even during obesity. This observation most likely reflects the female sex-associated robust induction of B-cell responses and humoral immunity, mediated by hormonal and chromosomal factors (56–58), which needs further exploration during obesity. It also suggests that greater disease severity in females is likely caused by a dysregulated innate immune response rather than a failure of humoral priming.

There are certain limitations in our study. We used only the mouse-adapted 2009 H1N1 IAV strain, and the results might vary with different strains and subtypes of influenza viruses. The females with obesity were smaller and developed less severe hyperglycemia than males with obesity. This might overestimate the interpretation of the biological sex difference. Estrous cycle variation in females could also modulate inflammatory responses and it could not be controlled in this model. The mouse model is inbred and may not entirely recapitulate the genetic diversity existing in humans. Though we showed the phenotypic difference, this study does not explain the underlying mechanisms, such as the role of specific sex steroids, adipokines, cytokines, and genetic differences between males and females with obesity that require further exploration in the future.

This study provides a temporal analysis of IAV pathogenesis between male and female mice, with and without obesity, following both the high- and low-dose IAV infection. Biological sex difference was observed in influenza severity, where females, irrespective of obesity status, developed more severe disease than males. Likewise, obesity-associated increase in disease severity was also observed in both male and female DIO mice, especially after the low-dose IAV infection. Females with obesity exhibited the greatest disease severity, likely driven by dysregulated inflammatory responses and disproportionate immune cell infiltration. These findings emphasize the need to account for biological sex differences in IAV pathogenesis studies and in the interpretation of preclinical, clinical, and epidemiological data in populations with obesity. It will be critical to better understand the mechanisms of immune dysfunction during virus infection in obesity and to develop safer and more effective preventive and therapeutic strategies.

## Materials and methods

### Animals and diet treatment

All animal procedures were carried out in accordance with the Institutional Animal Care and Use Committee-approved protocol (4855) at Kansas State University (KSU). Male and female C57BL/6J mice (strain 000664, Jackson Laboratory, Bar Harbor, ME, USA) were purchased at 4-5 weeks of age and after a week of acclimatization, randomly assigned either to a low-fat diet (LFD, 10 kCal%, D12450Ji, Research Diets, New Brunswick, NJ, USA) or a high-fat diet (HFD, 60 kCal%, D12492i, Research Diets, New Brunswick, NJ, USA) treatments. Each week, body mass was measured, and after 13 to 14 weeks, mice on HFD that gained ≥20% body mass compared to the average body mass of age- and sex-matched mice on LFD were defined as mice with obesity (59). Those on HFD but did not reach the obesity threshold were defined as non-responders.

### Glucose tolerance test (GTT)

GTT was performed on the 8^th^ and 13^th^ or 14^th^ weeks of diet treatment. For GTT, after 6 hours of fasting, a 25% glucose solution was administered intraperitoneally (dose: 2 g/Kg body mass). Blood glucose levels were measured at different time points, including before fasting (i.e., to measure the fed glucose), at 0 min (i.e., after fasting, and immediately before glucose administration to measure fasting glucose), and at 15, 30, 60, or 120 minutes after glucose administration using the AlphaTrak3 blood glucose monitoring system (Zoetis, NJ, USA) by pricking the tip of tail with a sterile lancet (59).

### Measurement of different metabolic biomarkers

Plasma concentrations of leptin (#90030, Crystal Chem, IL, USA); adiponectin (#80569, Crystal Chem, IL, USA); and total cholesterol (#STA-384; Cell Biolabs Inc., CA, USA) were measured following the manufacturer’s instructions, at week 0 (i.e., before diet treatment) and after 14 weeks.

### Virus infection and post-infection monitoring

The mouse-adapted A/California/04/2009 H1N1 IAVs grown in Madin-Darby Canine Kidney (MDCK) cells were used for virus infection after 13 or 14 weeks of diet treatment. For high-dose and low-dose infection studies, 10^3^ TCID_50_ and 10^1.5^ TCID_50_ doses were used, which in non-obese adult mice represent lethal and non-lethal doses, respectively (16). Viruses were diluted in Dulbecco’s Modified Eagle Medium (DMEM) and delivered intranasally (30 µL, 15 µL/nare) under the ketamine (80-100mg/Kg) and xylazine (5-10mg/Kg) anesthesia (16). DMEM-inoculated mice were used as uninfected controls for cytokine and chemokine analysis. During the day of inoculation, nose-to-anus length was also measured to determine BMI (i.e., body mass/height^2^). After infection, body mass and rectal temperature were recorded daily for 21 dpi to determine disease severity. Subsets of mice were euthanized at 3, 10, and 21 dpi to collect lung samples for virus titration, histopathology, flow cytometry, and cytokine and chemokine measurements. During the observation period, if any mouse lost ≥25% of its baseline body mass, had a temperature below 32^0^C, or showed severe respiratory signs, it was humanely euthanized (16).

### Virus titration

For virus titration, the MDCK cell-based 50% tissue culture infectious dose (TCID_50_) assay was performed (57). Briefly, 10-fold serially diluted lung homogenates were transferred to six replicates of MDCK cells in 96-well cell culture plates and incubated in a 32^0^C and 5% CO_2_ incubator. After 6 days of incubation, cells were fixed with 4% formaldehyde solution, stained with naphthol blue-black solution, and virus titers were determined using the Reed and Muench method.

### Antibody measurement

In-house standardized enzyme-linked immunosorbent assays (ELISAs) were used to measure IgG, IgG1, and IgG2c antibodies in the plasma and IgG and IgA antibodies in lung homogenates (56, 60). Plasma and lung homogenates were two-fold serially diluted starting with 1:250 and 1:20 dilutions, respectively. Horseradish peroxidase (HRP)-conjugated secondary antibodies used were IgG (#31430, Invitrogen, MA, USA), IgG1 (#PA1-74421, Invitrogen, MA, USA), IgG2c (#56970, Cell Signaling Technologies, MA, USA), and IgA (#626720, Invitrogen, MA, USA) (16).

### Histopathology

Left lung lobes were inflated with aqueous buffered zinc formalin fixative (Z-fix, #170, Anatech Ltd., MI, USA) on the day of necropsy and immersed in the fixative for a minimum of 72 hours before sending to the Kansas Veterinary Diagnostic Laboratory (KVDL) for histological processing. For comprehensive analysis, the left lung was sectioned into four quadrants, ensuring representative sampling across different anatomical regions. Fixed lung tissues were embedded in paraffin, sectioned at a thickness of 5 µm, mounted on glass slides, and stained with hematoxylin and eosin (H&E) for microscopic evaluation (60). A board-certified veterinary pathologist, blinded to the treatment groups, conducted a histopathological assessment. Inflammation of the pleura, parenchyma, vasculatures (arteries, veins, and capillaries), and airways (main bronchus, primary bronchi, and respiratory bronchioles) were each scored from 0 to 5. The cumulative lung inflammation score was between 0 to 40, reflecting the overall severity and extent of the inflammatory changes (59).

### Cytokines and chemokines measurement

Cytokines and chemokines in the lung homogenates were measured using either the ProcartaPlex™ mouse cytokine and chemokine Panel 1, 26plex (#EPX260-26088-901) or mouse immune monitoring panel, 48plex (#EPX480-20834-901, Thermo Fisher Scientific, MA, USA) (16). The 26-plex included IL-1 family cytokines (i.e., IL-1β and IL-18); common gamma chain family cytokines (i.e., IL-2, IL-4, and IL-9); TNFα; IL-10 family cytokines (i.e., IL-10 and IL-22); IL-12 family cytokines (i.e., IL-12p70, IL-23, and IL-27); IL-6, IL-17A, IL-5, and chemokines MCP-1, MIP-1α, GRO-α etc. Likewise, the 48-plex had interferons (IFNα, IFNγ, and IL-28 referred as IFN-λ); IL-1 family cytokines (i.e, IL-1α, IL-1β, IL-33, and IL-18); common gamma-chain cytokines (i.e., IL-2, IL-4, IL-7, IL-9, and IL-15); TNF superfamily cytokines (i.e., TNFα, RANKL, and BAFF); IL-10 family cytokines (i.e, IL-10, IL-19, IL-22); IL-12 family cytokines (i.e., IL-12p70, IL-23, and IL27); IL-6 family cytokines (i.e., IL-6, IL-31, and LIF); different chemokines (i.e., MCP-1, MIP-1α, MIP-1β, RANTES, MCP-3, Eotaxin, GRO-α, MIP-2α, CXCL-5, and IP-10); various growth factors (i.e., BTC, G-CSF, GM-CSF, M-CSF, IL-3, and VEGF-A); and Th-2 cytokines IL-5 and IL-13. Plate reading was carried out using Luminex xMAP technology (TX, USA). For analysis, raw data were imported into the ProcartaPlex analysis application (Thermo Fisher Scientific, Waltham, MA, USA), and cytokine concentrations (pg/mL) were determined using a five-parameter line fitted to the standards provided by the manufacturer. For samples not having measurable cytokine concentration, half of the lowest detected value was used, while for concentrations beyond the highest detection limit, 1.5 times the highest detection limit was used to enable statistical comparisons (16, 59).

### Preparation of cells for flow cytometry

Lungs were harvested, and single cells were prepared by enzymatic digestion (61). Briefly, lung lobes were chopped into fine pieces, lung digestion medium containing DNAse type I (diluted to 10^4^ U/mL, 8µL/sample, #04536282001, Roche, Basel, Switzerland) and Gibco™ collagenase II (diluted to 10 mg/mL, 100 µL/sample, #17101015, Thermo Fisher Scientific, MA, USA) were added, and tubes were incubated (37 °C and 5% CO_2_ incubator) with intermittent shaking at 10 min intervals for 40 minutes. After incubation, lung pieces were pressed through the Fisherbrand™ 70 µm cell strainer (#22363548, Thermo Fisher Scientific, MA, USA) followed by lysis of red blood cells using Gibco™ ACK lysis buffer (#A1049201, Thermo Fisher Scientific, MA, USA) (61).

### Flow cytometry

For staining with myeloid and lymphoid cell markers, 1 million cells were transferred into 96-well plates, Fc receptors were blocked using anti-CD16/32 antibody (#553141, BD Biosciences, CA, USA) and stained with panel-specific antibodies. Antibodies used in myeloid panels were BD Horizon V500 rat anti-mouse CD45 (#561487, clone 30-F11), BD Horizon BV786 mouse anti-mouse CD64 (#569507, clone X54-5/7.1), BD Horizon BV650 rat anti-CD11b (#563402, clone M1/70), BD Pharmingen PE-Cy7 hamster anti-mouse CD11c (#561022, clone HL3), BD Horizon RY610 rat anti-mouse Ly-6C (#571149, clone AL-21), BD Horizon BB700 rat anti-mouse Ly-6G (#566453, 1A8), BD Horizon BV421 rat anti-mouse CD86 (564198, GL1), BD Pharmingen PE rat anti-mouse Siglec-F (#562068, E50-2440), and BD Pharmingen Alexa Fluor 700 rat anti-mouse I-A/I-E (#570878, clone M5/114.15.2), BD Pharmingen Alexa Fluor 647 rat anti-mouse CD206 (#568808, clone Y17-505). Intracellular staining for CD206 was carried out following cell fixation with BD Cytofix fixation buffer (#554655) and permeabilization with BD Perm/Wash buffer (#554723).

Antibodies used in lymphoid panels were BD Horizon BV650 rat anti-mouse CD3 (#569683, clone 17A2), BD Pharmingen FITC rat anti-mouse CD4 (#561828, clone GK1.5), BD Horizon R718 rat anti-mouse CD8α (#566985, clone 53-6.7), BD Pharmingen PE-Cy7 rat anti-mouse CD19 (#561739, clone 1D3), BD Horizon BV421 mouse anti-mouse NK-1.1 (#562921, clone PK136), BD Horizon PE-CF594 hamster anti-mouse γδ T-cell receptor (#563532, clone GL3), BD Horizon RB705 rat anti-mouse CD25 (#570628, clone PC61), and BD Pharmingen PE rat anti-mouse Foxp3 (#563101, clone R16-715). For Foxp3 intracellular staining, cells were fixed and then permeabilized using BD Pharmingen Transcription Factor Buffer Set (#562574). Cells were acquired in LSR Fortessa X-20 (BD Biosciences, NJ, USA) and analyzed in FlowJo_v10.10.0_CL (BD Life Sciences, OR, USA).

From myeloid panel, frequencies of CD45^+^ cells, neutrophils (CD45^+^CD11b^+^Ly6G^+^), eosinophils (CD45^+^SiglecF^+^CD11c^-^), alveolar macrophages (CD45^+^SiglecF^+^CD11b^-/lo^CD11c^+^CD64^+^), interstitial macrophages (CD45^+^CD11b^+^MHC II^+^CD64^int/hi^), CD11b^+^ DC (CD45^+^CD11b^+^MHC II^+^CD64^-^), M1 macrophages (CD45^+^CD64^+^CD11b^+^CD86^+^), M2 macrophages (CD45^+^CD64^+^CD11b^+^CD206^+^), inflammatory (Ly6c^+^) monocytes/macrophages (CD45^+^CD64^+^CD11b^+^Ly6c^+^), and monocyte-derived dendritic cells (MODCs) (CD45^+^CD11b^+^MHCII^+/-^CD11c^+^Ly6c^+^) were determined (**Supplementary Figure 1**). Likewise, in lymphoid panels, frequencies of total T cells (CD3^+^), T helper cells (CD3^+^CD4^+^), CD8^+^ T cells (CD3^+^CD8^+^), regulatory T cells (CD3^+^CD4^+^CD25^+^Foxp3^+^), γδ T cells (CD3^+^γδ^+^), natural killer (NK) cells (CD3^-^NK1.1^+^), and B cells (CD3^-^B220^+^) were determined (**Supplementary Figure 2**).

### Statistical analysis

Data was analyzed using GraphPad Prism version 10.5.0 and Microsoft Excel for Microsoft 365 (version 2507). Normality test was performed using the Shapiro-Wilk test. Normally distributed data were compared using an unpaired t-test or one-way ANOVA followed by Tukey’s post-hoc analysis, while non-normally distributed data were analyzed by the Mann-Whitney test or Kruskal-Wallis test followed by Dunn’s post-hoc comparisons for two or more than two groups, respectively. If repeated measurements were taken from the same animal, they were compared using repeated measures ANOVA (mixed effects model) with Tukey’s multiple comparisons. For cytokine and chemokine analysis, unpaired t-tests followed by Holm-Sidak correction for multiple comparisons were used for two groups, and one-way ANOVA or Kruskal-Wallis test followed by Benjamini and Hochberg multiple comparisons were used. Data were considered statistically significant at p<0.05 and having a trend at 0.05≤p≤0.1.

## Acknowledgements

The mouse-adapted viruses were provided by Drs. Sabra Klein and Andrew Pekosz from Johns Hopkins Bloomberg School of Public Health. We acknowledge the Comparative Medicine Group (CMG) at Kansas State University (KSU) for their help during animal experiments. We also acknowledge the Animal Model/Pathology (AMP) and Molecular and Cellular Biology (MCB) cores of the Center on Emerging and Zoonotic Infectious Diseases (CEZID) at KSU for their help during animal protocol development and flow cytometry assays.

## Author contributions

Conceptualization, funding acquisition, project administration, and supervision: SD; Methodology, data curation, formal analysis, validation, visualization: SP, SV, BW, SBM, TA, SD; Writing – original draft: SP, SD; Writing – review and editing: SP, SV, BW, SBM, TA, SD.

## Data availability

All relevant data are readily available. While data supporting the results and conclusions are included within the paper, other related data will be provided in response to reasonable requests.

---

**Supplementary Figure 1:**
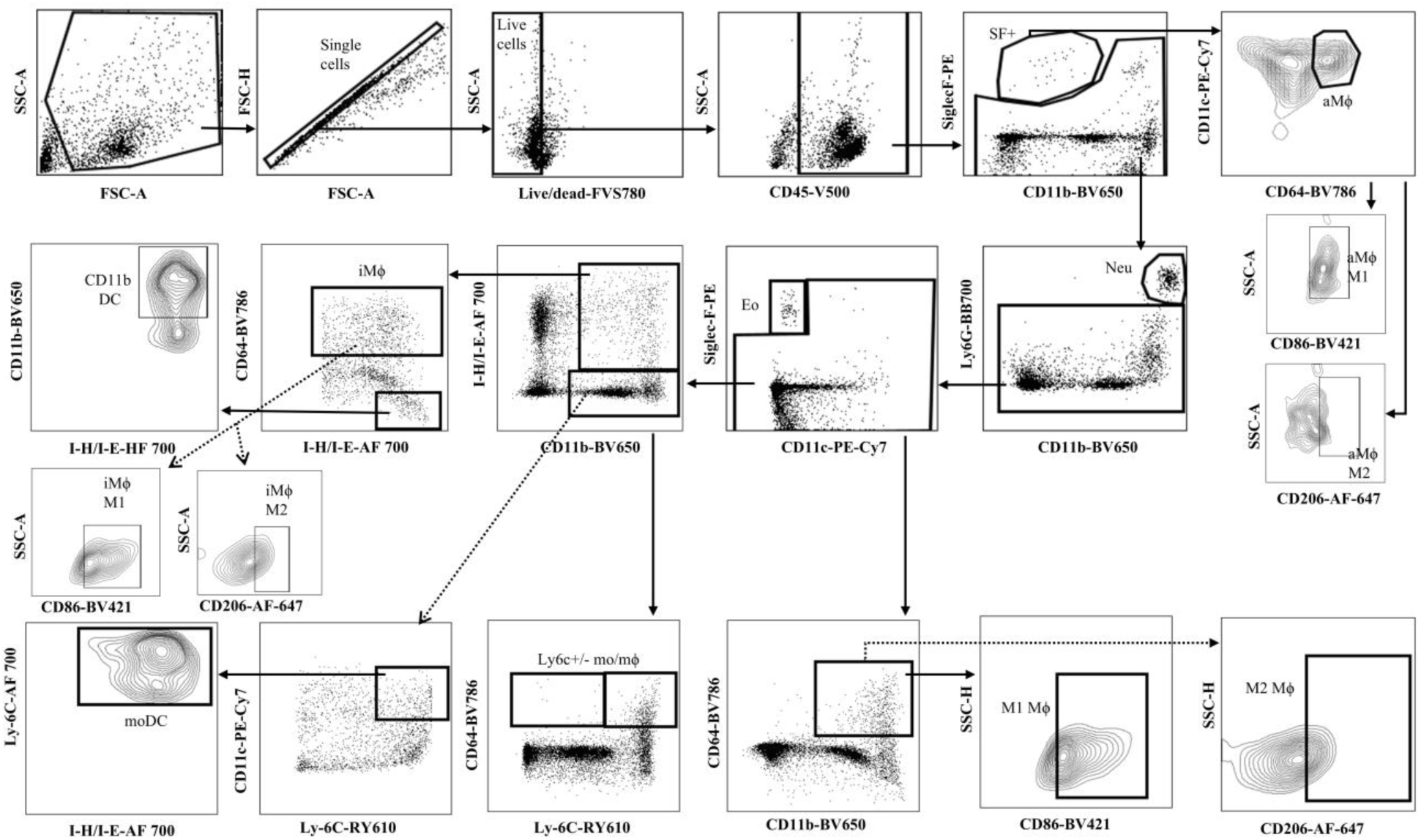
Gating strategy for myeloid cells in the lungs. Male and female C57BL/6J mice, with or without obesity, were infected with a low dose (i.e., 10^1.5^ TCID_50_) of the 2009 pandemic H1N1 virus. After 3 days post-infection (dpi), mice were euthanized, cells were prepared from the lungs, stained with different antibody markers, and acquired on a flow cytometer to determine the frequencies of different myeloid cell populations. The representative gating strategy is illustrated using data from one mouse. Total cells were gated, followed by single cells, and live cells using the live/dead staining dye. CD45^+^ cells were then selected and alveolar macrophages (CD45^+^SiglecF^+^CD11b^-/lo^CD11c^+^CD64^+^), neutrophils (CD45^+^CD11b^+^Ly6G^+^), eosinophils (CD45^+^SiglecF^+^CD11c^-^), interstitial macrophages (CD45^+^CD11b^+^MHC II^+^CD64^int/hi^), CD11b^+^ DC (CD45^+^CD11b^+^MHC II^+^CD64^-^), M1 macrophages (CD45^+^CD64^+^CD11b^+^CD86^+^), M2 macrophages (CD45^+^CD64^+^CD11b^+^CD206^+^), inflammatory (Ly6c^+^) monocytes/macrophages (CD45^+^CD64^+^CD11b^+^Ly6c^+^), and monocyte-derived dendritic cells (MODCs) (CD45^+^CD11b^+^MHCII^+/-^CD11c^+^Ly6c^+^) were gated.

**Supplementary Figure 2:**
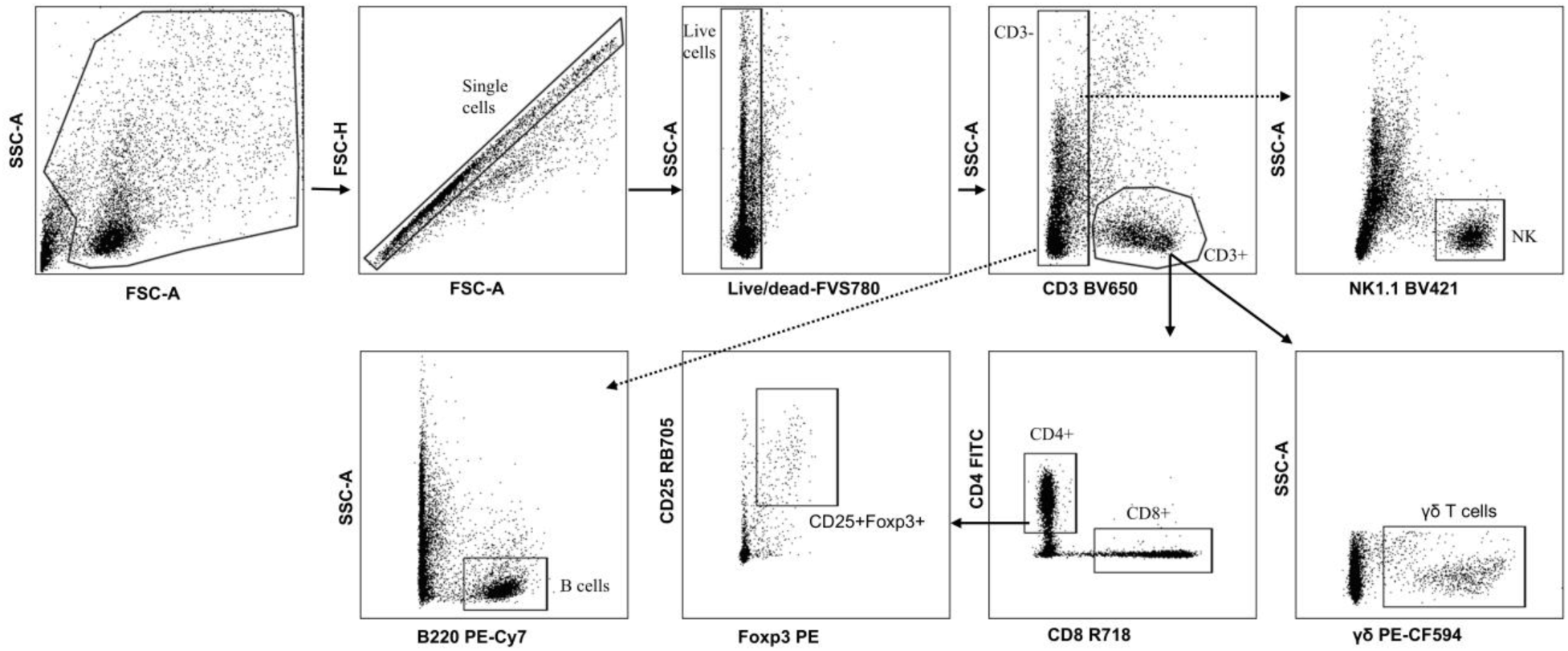
Gating strategy for lymphoid cells in the lungs. Male and female C57BL/6J mice, with or without obesity, were infected with a low dose (i.e., 10^1.5^ TCID_50_) of 2009 pandemic H1N1 virus. After 3 days post-infection (dpi), mice were euthanized, cells were prepared from the lungs, stained with different antibody markers, and acquired on flow cytometer to determine the frequencies of different lymphoid cell populations. The representative gating strategy is illustrated using data from one mouse. Total cells were gated, followed by single cells, and live cells using the live/dead staining dye. CD3^+^ cells were gated followed by gating for T helper (CD3^+^CD4^+^) and CD8^+^ T cells (CD3^+^CD8^+^). CD4^+^ cells were further gated to determine the frequency of regulatory T (Treg) cells (CD3^+^CD4^+^CD25^+^Foxp3^+^). CD3^-^ cells were used to determine the frequencies of NK cells (CD3^-^NK1.1^+^), gamma-delta T cells (CD3^-^γδ^+^), and B cells (CD3-B220+).

**Supplementary Figure 3.**
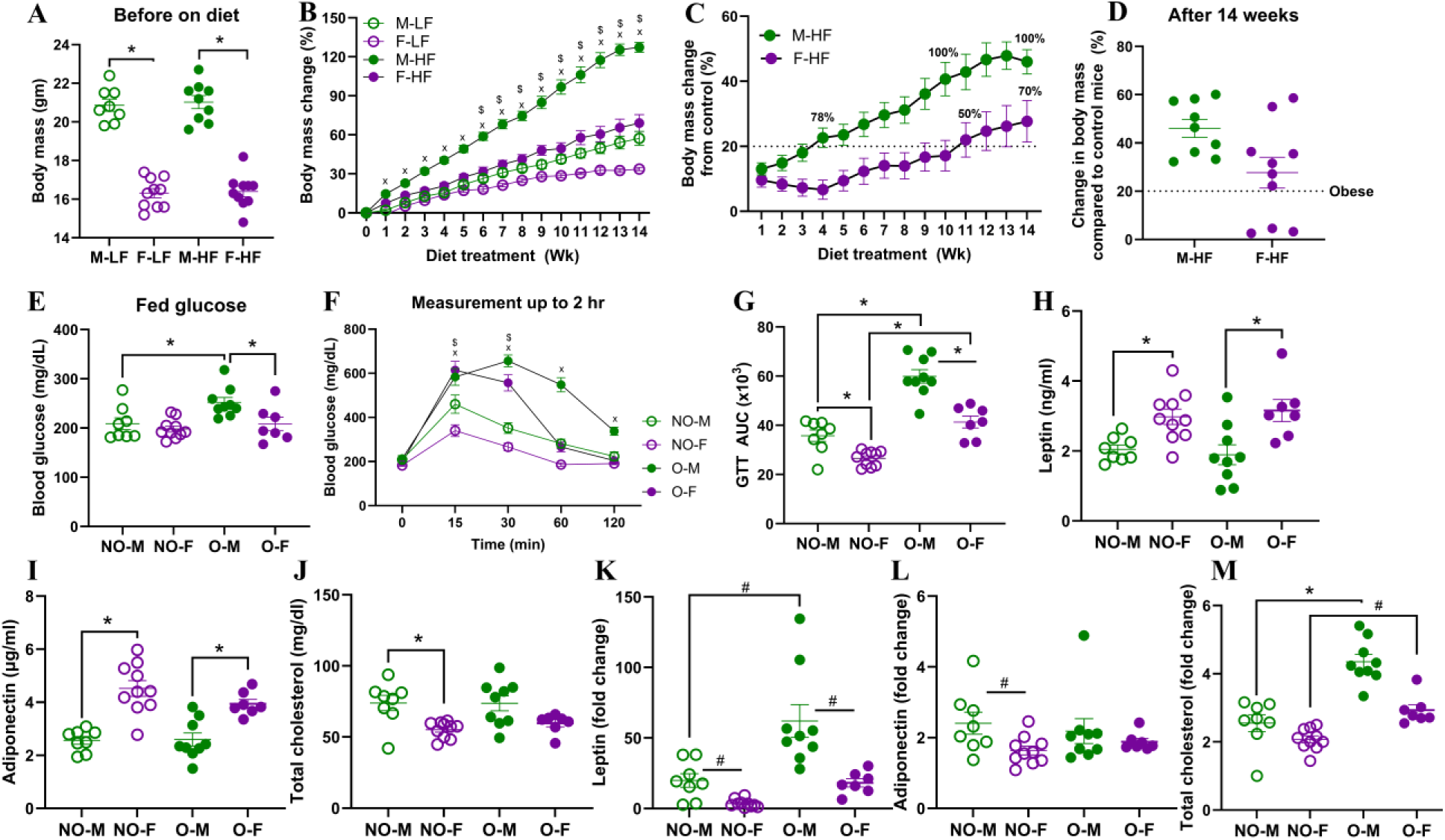
Body mass and blood parameters after diet treatment in male and female mice. C57BL/6J male and female mice of age 5-6 weeks were treated with a low-fat (LF) diet or a high-fat (HF) diet for 14 weeks. (A) Body mass before the diet treatment started; (B) percentage change in body mass following the diet treatment; (C) percentage change in body mass from age- and sex-matched controls over time, and (D) at the 14^th^ week are shown. The Glucose tolerance test (GTT) was carried out on the 8^th^ week of diet treatment, and data of mice that ultimately became obese or non-obese were compared. (E) Fed glucose; (F) GTT over 2 hours; and (G) GTT area under the curve (AUC). Blood biomarkers were also measured before diet treatment and after the 14^th^ week of diet treatment. The concentrations of (H) leptin, (I) adiponectin, and (J) total cholesterol before diet treatment, and (K-M) fold changes in these biomarkers by the end of the 14^th^ week, respectively. Data is shown as mean ± standard error of the mean (SEM) (n=7-10/group). Statistical comparisons were made using one-way ANOVA and Tukey’s test or Kruskal-Wallis and Dunn’s post-hoc tests. Data in B and F were measured using two-way repeated measures ANOVA (mixed model) followed by Tukey’s multiple comparisons. An asterisk (*) indicates a statistically significant difference (p<0.05) while a hash (^#^) indicates a trend (0.05≤p≤0.1). In figures B and F, ‘x’ and ‘$’ represent significant differences between non-obese (or LF) versus obese (or HF) males and females, respectively. Abbreviations: M-LF, males on low-fat diet; F-LF, females on low-fat diet; M-HF, males on high-fat diet; F-HF, females on high-fat diet; NO-M, non-obese males; NO-F, non-obese females; O-M, males with obesity; and O-F, females with obesity.

**Supplementary Figure 4.**
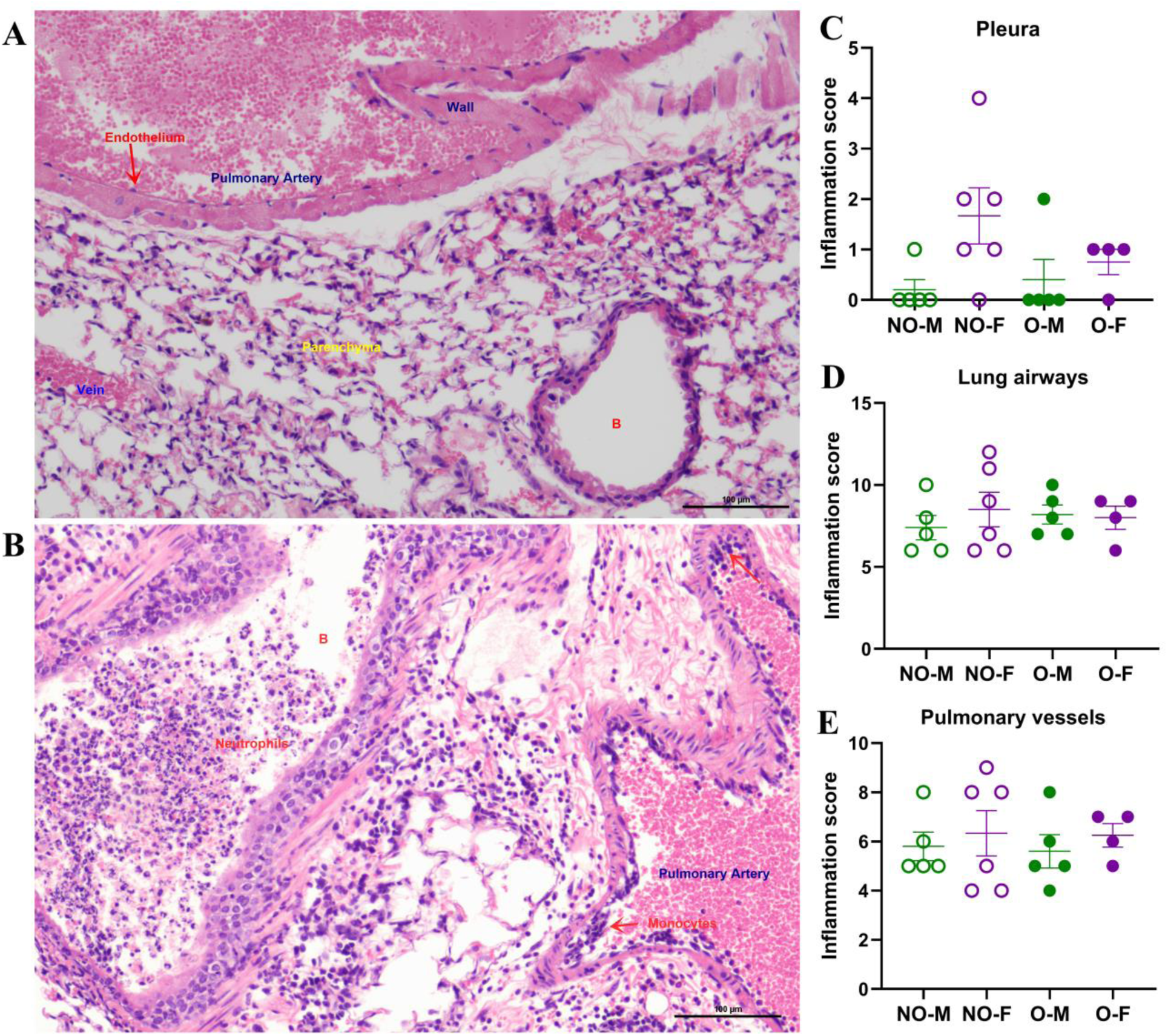
Inflammatory changes in the lungs after a high-dose infection. Males and females, with or without obesity, were infected with a high dose (i.e., 10^3^ TCID_50_) of the 2009 pandemic IAV, and at 3 days post-infection, lungs were fixed and used for histopathological analysis using H&E staining. Representative lung images from (A) medium-inoculated and (B) virus-inoculated mice are shown. Inflammatory changes in the (C) pleura, (D) lung airways, and (E) pulmonary vessels are compared. Data is shown as mean ± standard error of the mean (SEM) (n=5-6/group). Statistical comparisons were made using one-way ANOVA and Tukey’s post-hoc test or Kruskal-Wallis and Dunn’s multiple comparisons. Abbreviations: NO-M, non-obese males; NO-F, non-obese females; O-M, males with obesity; and O-F, females with obesity.

**Supplementary Figure 5.**
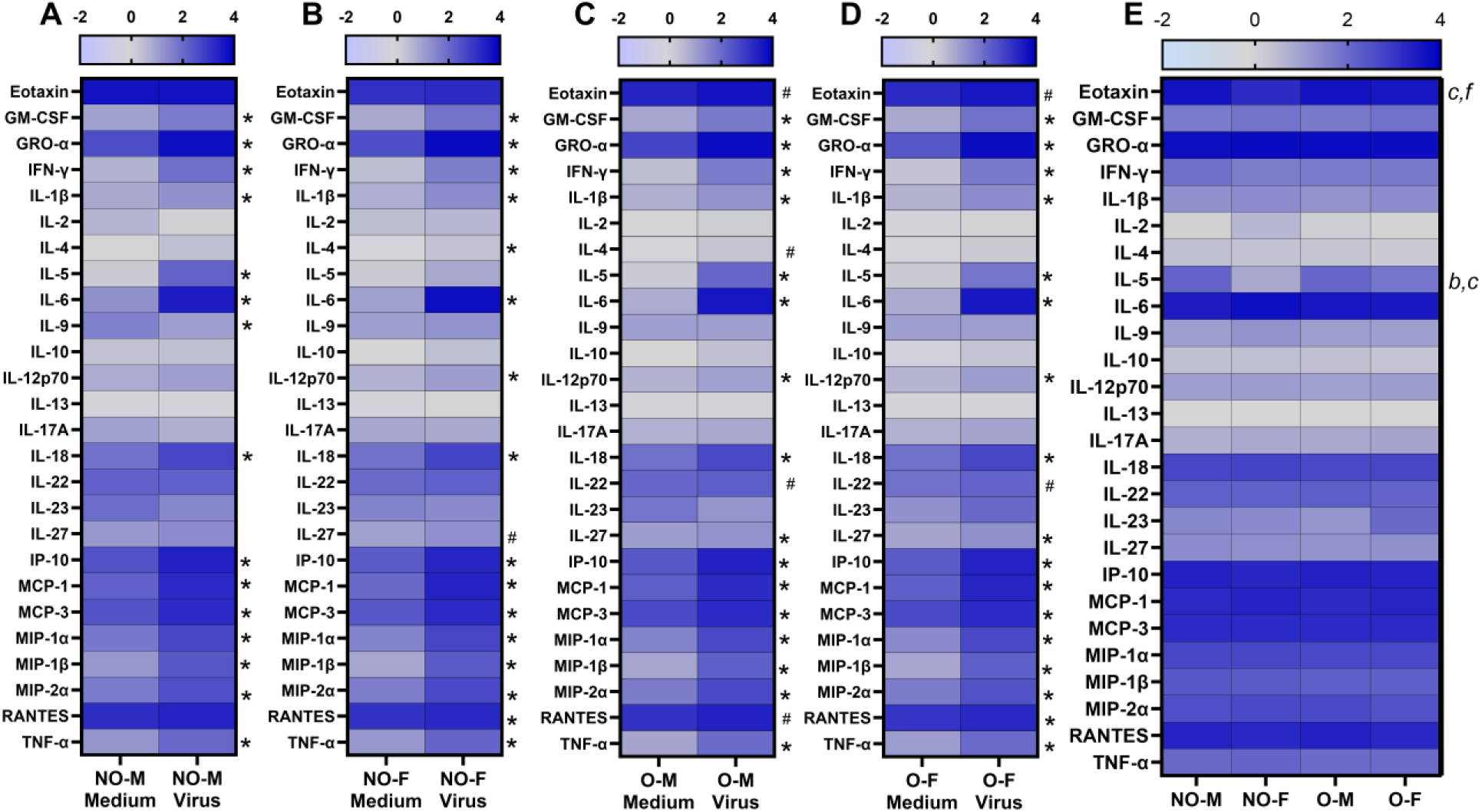
Cytokines and chemokines in the lungs following high-dose IAV infection. Males and females, with or without obesity, were infected with a high dose (i.e., 10^3^ TCID_50_) of the 2009 pandemic H1N1 virus or vehicle (i.e., medium) only. At 3 days post-infection (dpi), mice were euthanized, and cytokine and chemokine responses were measured in the lung homogenates. Comparison of absolute concentrations of log_10_-transformed cytokine and chemokine responses between medium-inoculated and virus-infected (A) non-obese males (NO-M), (B) non-obese females (NO-F), (C) obese males (O-M), and (D) obese females (n=3-6/group). Likewise, (E) comparisons of cytokines and chemokines among virus-infected NO-M, NO-F, O-M, and O-F are shown. Statistical analysis was carried out by unpaired t-tests, followed by Holm-Sidak correction for multiple comparisons, and one-way ANOVA or Kruskal-Wallis test, followed by Benjamini and Hochberg multiple comparisons. Data were considered statistically significant at p<0.05 and having a trend at 0.05≤p≤0.1. Figures A-D: an asterisk (*) represents a significant difference, and a hash represents a trend. In Figure E, a and e; b and f; c and g; and d and h represented a significant difference or a trend between NO-M versus O-M; NO-F versus O-F; NO-M versus NO-F; and O-M versus O-F, respectively.

**Supplementary Figure 6.**
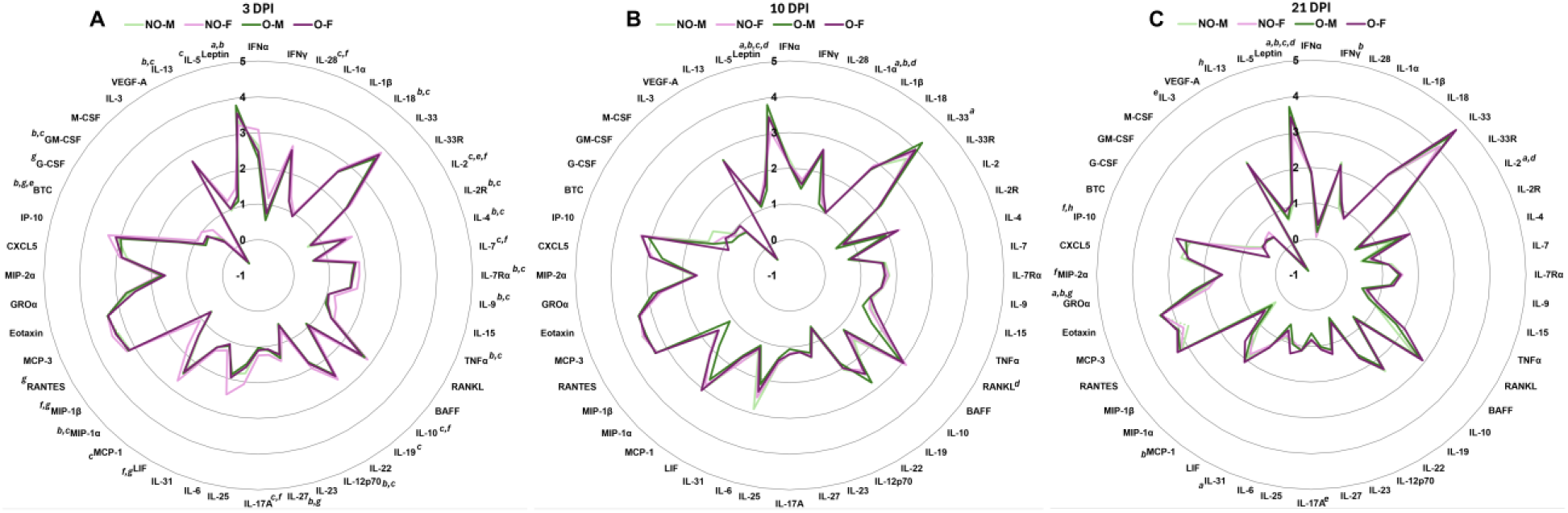
Cytokine and chemokine responses in the lungs following low-dose IAV infection. After infection with a low dose (i.e., 10^1.5^ TCID_50_) of the 2009 pandemic H1N1 IAV, subsets of mice were euthanized at 3-, 10-, and 21-days post-infection (dpi), lungs were collected, and various cytokines and chemokines were measured. Absolute concentrations of cytokines and chemokines (log_10_-transformed) among virus-infected mice euthanized at (A) 3 dpi, (B) 10 dpi, and (C) 21 dpi are compared. Statistical comparison was made using one-way ANOVA or Kruskal-Wallis test, followed by Benjamini and Hochberg multiple comparisons. Data were considered statistically significant at p<0.05 and having a trend at 0.05≤p≤0.1. Letters a and e; b and f; c and g; and d and h represented a significant difference or a trend between non-obese males (NO-M) versus obese males (O-M); non-obese females (NO-F) versus obese females (O-F); non-obese males (NO-M) versus non-obese females (NO-F); and obese males (O-M) versus obese females (O-F), respectively.

